# Characterization of Integrase and Excisionase Activity in Cell-free Protein Expression System Using a Modeling and Analysis Pipeline

**DOI:** 10.1101/2022.10.05.511053

**Authors:** Ayush Pandey, Makena L. Rodriguez, William Poole, Richard M. Murray

## Abstract

We present a full-stack modeling, analysis, and parameter identification pipeline to guide the modeling and design of biological systems starting from specifications to circuit implementations and parameterizations. We demonstrate this pipeline by characterizing the integrase and excisionase activity in cell-free protein expression system. We build on existing Python tools — BioCRNpyler, AutoReduce, and Bioscrape — to create this pipeline. For enzyme-mediated DNA recombination in cell-free system, we create detailed chemical reaction network models from simple high-level descriptions of the biological circuits and their context using BioCRNpyler. We use Bioscrape to show that the output of the detailed model is sensitive to many parameters. However, parameter identification is infeasible for this high-dimensional model, hence, we use AutoReduce to automatically obtain reduced models that have fewer parameters. This results in a hierarchy of reduced models under different assumptions to finally arrive at a minimal ODE model for each circuit. Then, we run sensitivity analysis-guided Bayesian inference using Bioscrape for each circuit to identify the model parameters. This process allows us to quantify integrase and excisionase activity in cell extracts enabling complex-circuit designs that depend on accurate control over protein expression levels through DNA recombination. The automated pipeline presented in this paper opens up a new approach to complex circuit design, modeling, reduction, and parameterization.

## Introduction

Over the past few years, there has been widespread adoption of software tools in synthetic biology research for modeling, simulation, analysis, data exchange, and design optimizations. The focus on bio-design automation and rational design in synthetic biology has led to this enthusiastic acceptance of software tools. A few examples include Synthetic Biology Open Language (SBOL)/Systems Biology Markup Language (SBML) compatible tools (for data and model standardization),^1,2^ COPASI^3^ (for modeling and simulation), iBioSim^4^ (for CAD-style circuit modeling), Tellurium^5^ (for text scripting of circuit models), promoter/RBS calculators^6,7^ (for prediction of transcription/translation initiation rates), and automated design recommendation tools.^8,9^

The rise in tools for specific tasks has led to their integration into pipelines and repositories like SynBioHub,^10^ Galaxy SynBioCAD,^11^ and Infobiotics.^12^ SynBioHub is a platform that facilitates the integration of software and data for synthetic biological designs so that users can easily share and reproduce system designs in a standardized format. Similarly, an end-to-end metabolic design portal called Galaxy SynBioCAD chains tools together into various workflows for common design and analysis tasks such as genetic design, flux balance analysis, and pathway benchmarking. While SynBioHub is focused on the standardization of software and data used in synthetic biology pipelines, Galaxy SynBioCAD allows for automated engineering and analysis workflows for designing metabolic pathways. A similar experimental design automation software that focuses on the test and learn parts of the design-build-test-learn cycle called RoundTrip^13^ has been developed recently. An important aspect of rational synthetic biology design that is not addressed by these approaches is model-guided design, or forward engineering, where specifications of circuits can be converted to mathematical models with parameters inferred from experimental data. A software suite that claims to integrate modeling, simulation, and verification for synthetic biological circuits was recently presented, called the Infobiotics Workbench.^12^ Infobiotics develops a domain-specific language for synthetic biology circuits to write specifications and provides a Java-based GUI platform for modeling and analyses. It is clear from these recent efforts that, for scalable biological circuit design, there is a need for open-source, user-friendly, and easy-to-access modeling and analysis pipelines that work alongside experimental data in a design-build-test-learn cycle.

Towards that end, we present an automated Python pipeline for iterative modeling, model reduction, analysis, and parameter identification of synthetic biological circuits. We further develop the Python software packages — BioCRNpyler^14^ (to build models), AutoReduce^15^ (to obtain reduced models), and Bioscrape^16^ (for simulations, analysis, and Bayesian inference using Emcee^17^) — to create the computational framework shown in Figure 1. A key idea with the proposed pipeline is its iterative nature by breaking down the system analysis into smaller parts. Once we learn the parameters of subsystem #1 and the system context, we fix these parameters for the next iteration. In this way, the parameterization of the model with both subsystem #1 and #2 is feasible and leads to reliable predictions that can guide the experimental design for the bigger system. This process of system identification and learning by parts to guide the design of more complex circuits can be extremely important for scalable biological circuit design and analysis. The proposed pipeline is a step in this direction. Further, the pipeline uses standardized data and model formats to allow for interoperability of tools and integration with other existing pipelines. To demonstrate the tools, we apply this pipeline to characterize an integrase and excisionase-mediated DNA recombination circuit in a cell-free extract.

**Figure 1:**
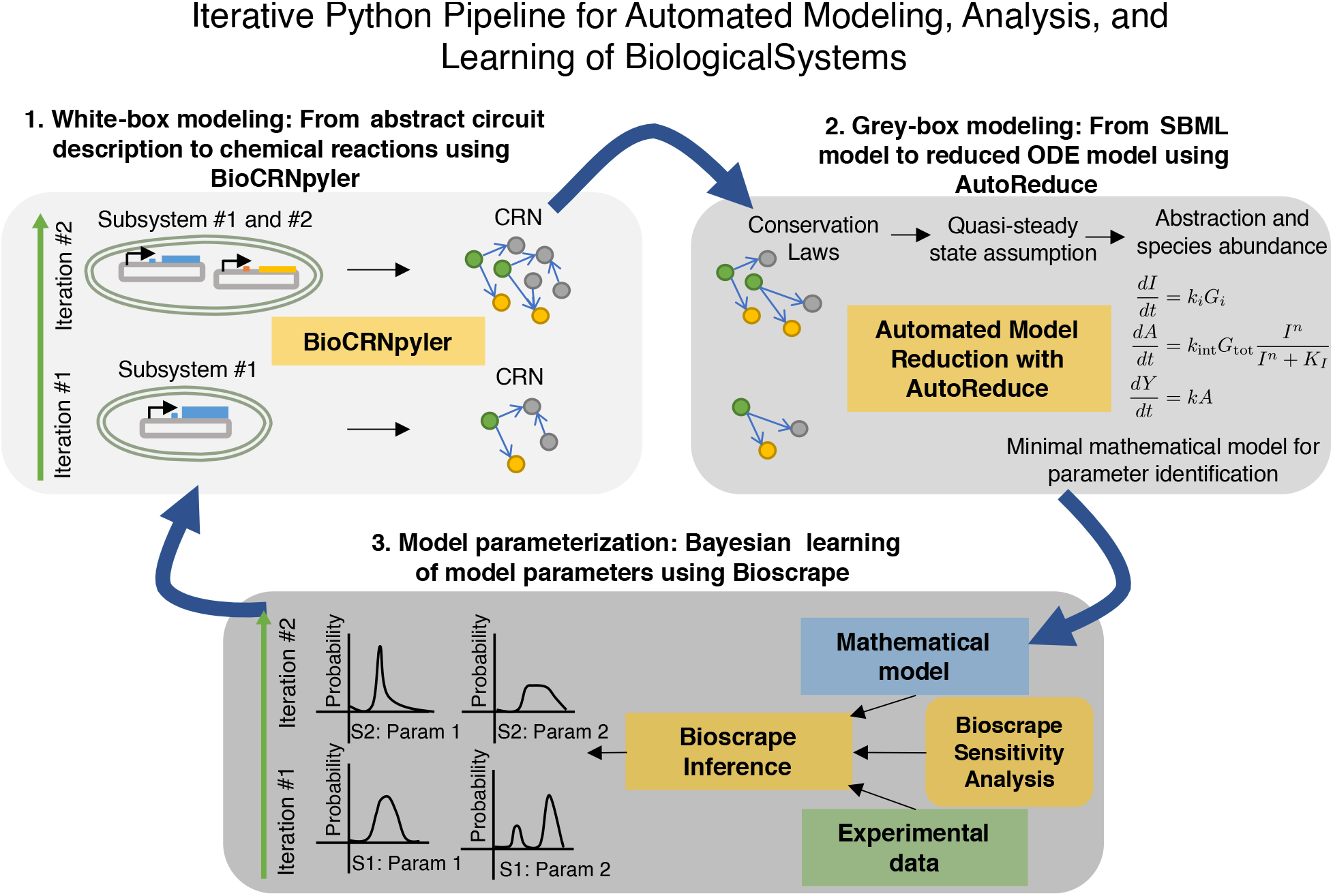
An iterative Python pipeline for modeling, analysis, and learning of biological circuits. In the first iteration, we build the CRN model for subsystem #1 and then obtain the minimal representation suitable for parameter identification. Bayesian inference is used to find parameter distributions. The results of the Bayesian inference are fed back into the more detailed model that is built in the second iteration. The previously identified context and circuit parameters are fixed in this larger model so that the analysis can now focus on the new parts introduced in the circuit. For each iteration, a CRN model is compiled using BioCRNpyler, this is the white-box modeling step that includes the mechanistic details of the system. With AutoReduce, we obtain a hierarchy of lower dimensional ODE models under different modeling assumptions. We call this grey-box modeling because we can tune the granularity of the models. Finally, we validate the model parameters in the reduced model by using Bioscrape.

### DNA Recombination Circuits

Recombinase-based circuits are circuits that utilize the unique functionality of phage integrases. Phage integrases are responsible for catalyzing the site-specific insertion of bacteriophage DNA into a host genome. These enzymes utilize specific attachment sites found in both the phage and host DNA: attP and attB, respectively. Upon integration of the phage DNA into the host genome, the attachment sites attL and attR are formed .^18^ This insertion can then be reversed by the expression of a similar enzyme called, excisionase. Excisionases work in conjunction with the integrases to excise the phage DNA from the host DNA, restoring the original attB and attP sites.^19^ Further research and characterization of these enzymes led to the discovery that the presence of anti-parallel attB and attP sites in the same strand of DNA leads to the inversion of the flanked segment, generating anti-parallel attL and attR sites. This attL and attR flanked DNA segment can then be reverted to its original orientation by an integrase-excisionase complex.^20^

One class of these phage integrases is serine integrases. Serine integrases have been used in many recombinase-based circuits for some of their advantageous characteristics when compared to other integrases.^21–23^ For example, some serine integrases only require an integrase and a reverse directionality factor, or excisionase, to function as opposed to other integrases which require additional cofactors for proper functioning. Moreover, serine integrases can use relatively short attachment sites when compared to other classes of integrases.

To date, recombinase-based circuits with various functionality and applications have been designed and experimentally validated. For example, event ordering detection circuits,^24^ gene networks that count,^25^ boolean logic and memory circuits,^26^ and many more circuits have all been created in living cells using recombinases. Although there has been progress in exploring the mechanisms for enzyme-mediated DNA recombination, the dynamics and characteristics of integrases and excisionases have not yet been fully quantified. In order to accelerate the design and implementation of increasingly complex recombinase circuits, novel methods are required to efficiently quantify their complex dynamics.

We used the proposed modeling and analysis pipeline to study the dynamical models of integrases and excisionases. We anticipate our pipeline providing a foundation to guide further design and implementation of increasingly complex recombinase circuits. In applying this pipeline, we built mathematical models and validated the mechanistic parameters for Bxb1 – a well-characterized and studied serine integrase-excisionase system.^27–29^ This pipeline, however, can be used to characterize general biological systems to guide the design of progressively complex circuits.

### Main Contributions

1. We develop new chemical reaction network (CRN) mechanisms in BioCRNpyler for integrase action to flip a promoter and the excisionase action to reverse it. These mechanisms model the details of the intermediate complexes and the binding rates of integrase to the DNA, excisionase to the integrase, the complexes, and to the DNA. With model reduction and *in silico* analysis, we show which mechanisms describe the expected behavior and then validate these models with *in vitro* cell-free data.
2. We further develop the Python model reduction package, AutoReduce, by providing new model reduction methods for CRN models and seamless import and export of models as SBML files.
3. For parameter inference, we extend the Bioscrape package by developing an easy-to-use parameter identification wrapper around Python emcee software. Although we only demonstrate time-series data in this paper, the Bioscrape inference package can also be used to validate models using distributional data from flow cytometry, and end-point data under multiple conditions.
4. We show a successful implementation and experimental validation of integrase and excisionase mediated DNA recombination circuit in cell-free system. After compiling, reducing, and parameterizing our model from experimental data, we are able to accurately model the effect of integrase and excisionase gene concentrations on fluorescent reporter output. For example, on adding higher integrase DNA concentrations, our models correctly predict the experimental observation that the reporter expression increases while it decreases as more excisionase is expressed.

## Results and Discussion

### Modeling Integrase Activity in Cell-free Systems

To characterize the integrase activity independent of the excisionase, we design a two plasmid system — (1) Bxb1 integrase expressing plasmid fused with CFP to measure integrase expression, and (2) a YFP plasmid that gets activated on integrase action (shown in Figure 2). Using this circuit, we characterize the integrase expression in a cell-free extract and its flipping action on a promoter to control the fluorescent reporter expression (data shown in Figure 5A, also see information on cell-free extracts in the Methods section).

**Figure 2:**
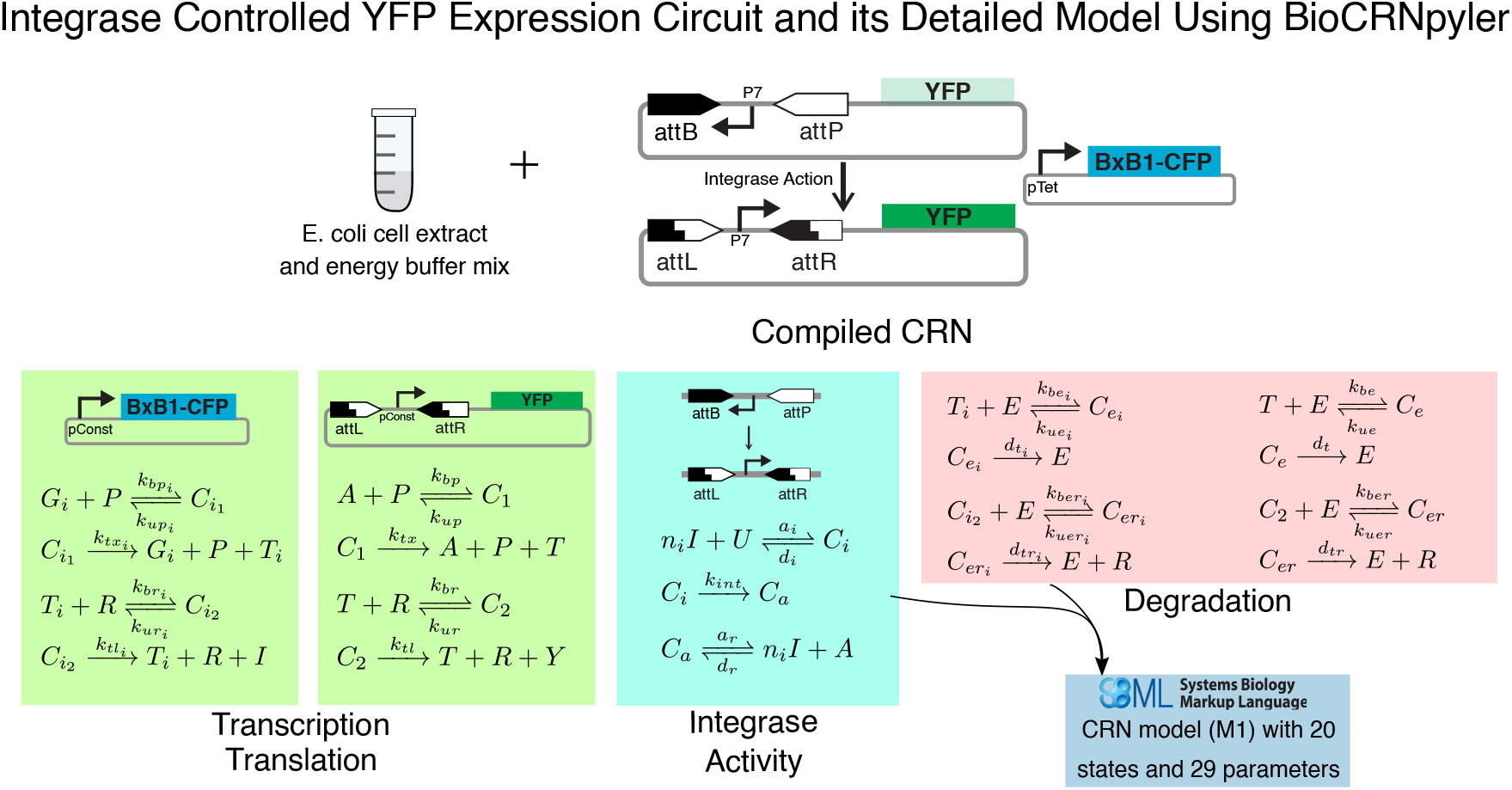
Modeling of integrase expression and activity in cell-free systems. The circuit consists of two plasmids, one expressing the Bxb1 integrase and the other with a reversed promoter upstream of the YFP reporter. The integrase activity flips the promoter so that YFP is expressed. BioCRNpyler is used to convert this abstract system description into a CRN model written as an SBML file with 20 species and 29 parameters. All species definitions in the model are given in the model details section in the supplementary information.

Towards this first goal, we model the two plasmid system in a cell-free extract using a detailed mechanistic chemical reaction network (CRN) with mass-action kinetics using BioCRNpyler. In BioCRNpyler, a CRN is compiled by specifying the circuit parts as `Component` objects, their interactions as `Mechanism` objects, and the context for the circuit as a `Mixture`. For the integrase circuit, we create an integrase component and two new mechanisms — a simple integrase flipper mechanism with one reaction that converts the inactive DNA to active by flipping the promoter direction and a detailed integrase flipper mechanism that models the binding reactions of the integrase to various DNA sites. Finally, to compile a CRN model, we use an existing BioCRNpyler mixture, `TxTlMixture` to model the cell-free context and resources. The detailed reactions for this model are given in Figure 5A. We simulate this CRN model using Bioscrape (shown in Figure 3) to explore the design space and the resource-loading effects. However, the detailed model is infeasible to fit the experimental data due to the problem of unidentifiability^30^ and high-dimensionality. Hence, we use AutoReduce to automatically derive potential reduced models for this system (see model reduction section in Methods). First, we derive and apply the conservation laws to reduce the system model. Then, we apply quasi-steady state approximation (QSSA) to obtain reduced models under different assumptions. Using model reduction performance metrics, we choose reduced models that recover the desired properties (integrase flipping, fluorescent reporter levels, and any other important context effects), shown in Figure 4 as M-3. See the performance metrics section in Methods to read more about how we assess reduced model performance. To obtain a further reduced model, we abstract the context by switching off resource-dependent mechanisms for transcription and translation in BioCRNpyler (more information on context abstraction is given in the Methods section). Then, we further reduce the one-step transcription and translation model using QSSA and assuming the abundance of certain species to obtain a minimal ODE model (M-4). It is evident from Figure 4 that the model M-4 recovers the commonly used Hill function, however, no heuristics were used in deriving this model. We export this model as SBML for further analysis.

**Figure 3:**
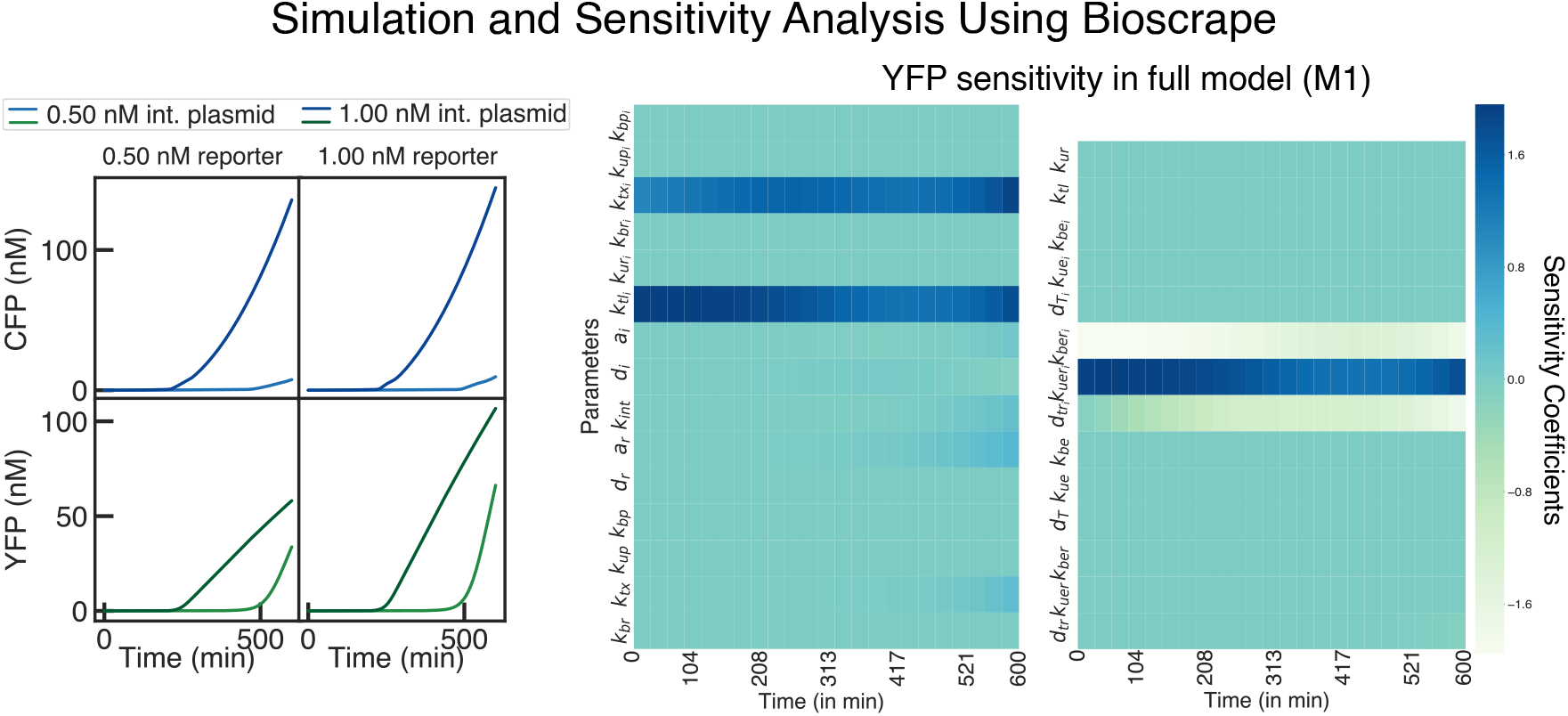
Simulation and analysis of integrase expression and activity in cell-free systems using Bioscrape. The simulations predict the CFP and YFP expressions under different initial conditions. The sensitivity analysis shows the most influential parameters for the time course of YFP expression.

**Figure 4:**
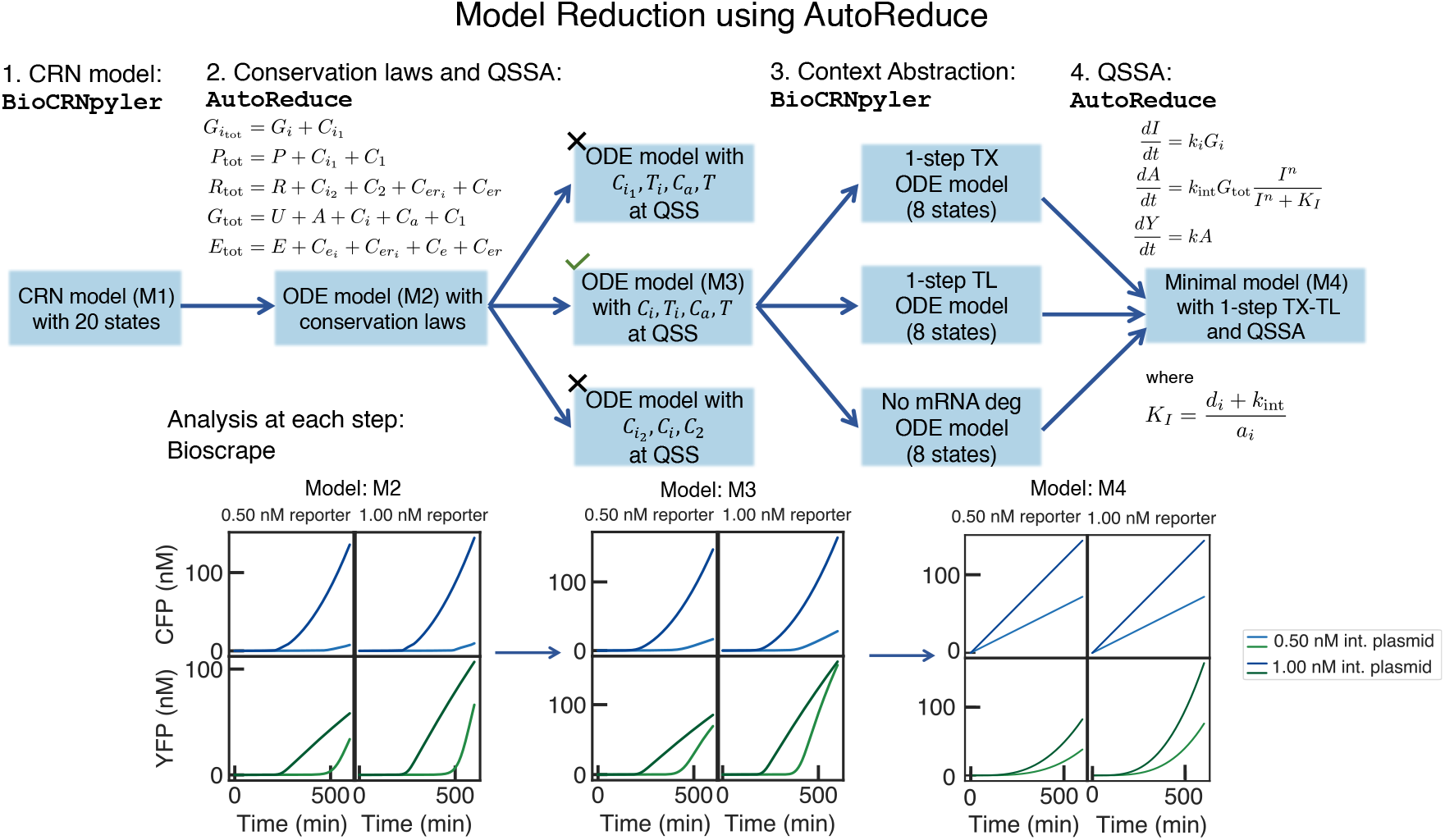
A hierarchy of reduced models for integrase expression and activity in cell-free systems using AutoReduce. The CRN model is reduced in multiple steps with AutoReduce. First, the conservation laws are determined (as shown in C-2) and a reduced model is obtained symbolically. This reduced model has 5 fewer states than the full model and is further reduced using QSSA. Multiple reduced models are possible at this stage, out of which one model, M3, is selected (tick marked in the figure) based on the error performance metric as computed by AutoReduce. Then, a minimal model is obtained by abstracting the details and further reducing the model using QSSA and species abundance assumptions. The minimal ODE model (M4) is shown in C-4. The simulations for reduced models (M2, M3, and M4) are also shown.

At each step in the model reduction hierarchy, the number of states and the parameters are reduced. For better accuracy of the reduced model, the user may choose a model that is higher up in the hierarchy, whereas, for faster simulations, the minimal model with the fewest parameters may be chosen. The computational run time to obtain each reduced model varies from a few seconds up to a maximum of a few minutes on an i7-6700K Intel CPU laptop with 16GB of RAM.

For the minimal model, we find the identifiable parameters as the most sensitive parameters affecting the measurements. The sensitivity analysis tools in Bioscrape (shown in Figure 5B) show the sensitivity of each model parameter with time for each output measurement (CFP and YFP). For the identifiable parameters, we use Bayesian inference tools in Bioscrape to fit the cell-free data (see Figure 5B and the inference section in Methods). The parameter inference algorithm is implemented in Bioscrape as a black-box Python wrapper for the emcee package that implements a Markov Chain Monte Carlo (MCMC) sampler for Bayesian inference. We import the SBML file for the model and the experimental data as CSV and then run the inference after choosing appropriate MCMC parameters (see Supplementary Information). After running the sampler, we obtain posterior distributions for the parameters which are then used for plotting the identified model simulation against the experimental data. Hence, the “full-stack” Python pipeline of modeling, design-space exploration, sensitivity analysis, model reduction, and parameter inference gives us a validated mathematical model for cell-free integrase activity.

**Figure 5:**
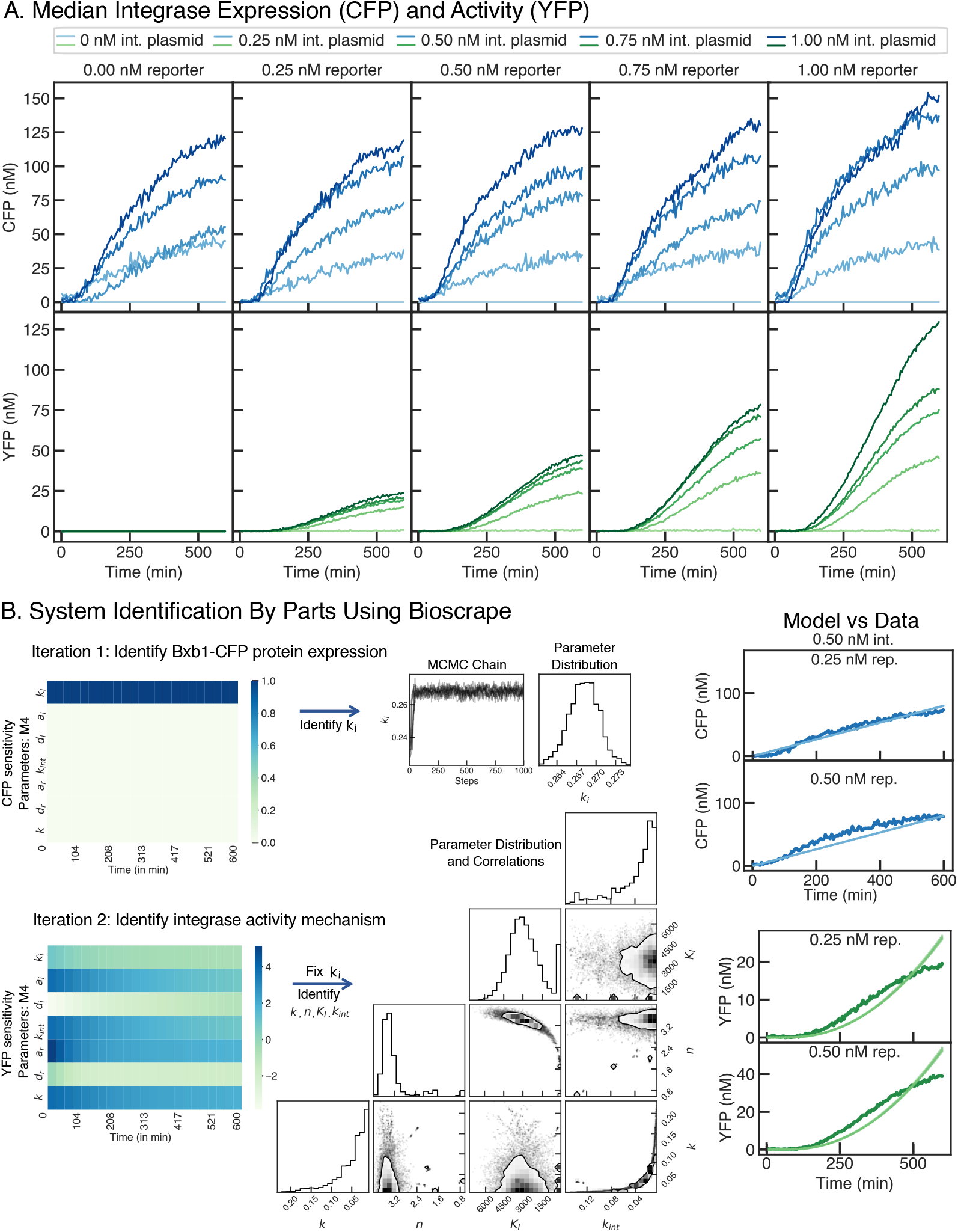
Experimental data and system identification by parts of cell-free integrase expression and activity. (A) Median background-subtracted fluorescence data for the integrase circuit in the cell-free extract. (B) We identify the system by parts, that is, we first select the integrase expression part of the circuit and run sensitivity analysis to find out its identifiable parameters. We observe that *k_i_* is the only sensitive parameter and hence, we run Bayesian inference to identify the posterior parameter distribution for *k_i_*. The model fit alongside the data is shown in the rightmost column. Once we have identified this part, we fix the corresponding parameter, *k_i_*, and run the sensitivity analysis for YFP output. We identify all parameters that YFP is sensitive to. The corner plot shows the posterior distributions of each parameter alongside their correlations with 75% confidence contours. The mismatch in the data and the model is due to the minimal model not capturing the plateauing expression as cell-free extract stops protein expression.

### Modeling Excisionase Activity in Cell-free Systems

Integrase action activates the output fluorescence protein expression by recombining the attP-attB site on the DNA to the attL-attR site so that the promoter is flipped towards the protein coding sequence. To accurately control this expression, we use the reverse directionality factor or the excisionase enzyme which can reverse the promoter direction by changing from the attL-attR site back to the “inactive” attP-attB site on the DNA. We design a new plasmid that expresses the excisionase fused with mScarlet to measure the excisionase expression (see Figure 6).

**Figure 6:**
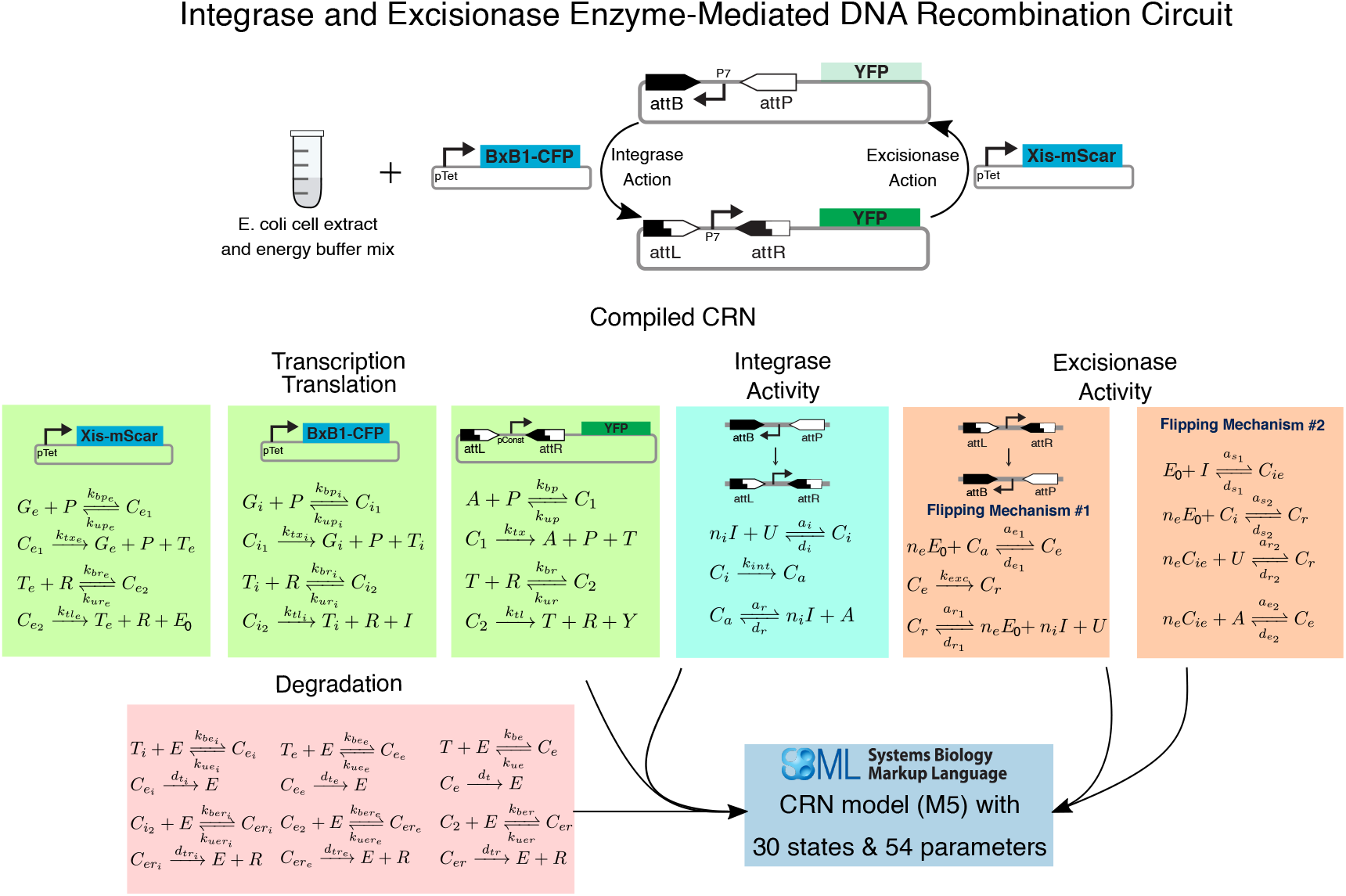
CRN models for excisionase expression and activity in cell-free extract using BioCRNpyler. We obtain a CRN for the circuit with both integrase and excisionase in cell-free extract by describing the circuit specifications in BioCRNpyler. The resulting model has 30 states and 54 parameters. The species definitions are given in the supplementary information on model species.

We build on the integrase models (both detailed and simplified mechanisms) shown in the previous section to develop excisionase mechanisms. We hypothesize that the excisionase can have two possible mechanisms to reverse the integrase activity:

1. The excisionase binds to the integrase and the resulting complex then binds to the attL-attR site on the DNA to flip it to the attP-attB site. The integrase-excisionase complex may also bind to the attP-attB site, sequestering the site from further recombinations.
2. The excisionase binds to the integrase-bound DNA sites to form a complex that then flips the promoter region or keeps it sequestered.

With BioCRNpyler, it is possible to switch ON any one or both of these mechanisms when compiling the CRN model. See Figure 6 for the list of chemical reactions. We compile a detailed integrase-excisionase model in BioCRNpyler with these mechanisms added into the same `TxTlMixture` as was used for the integrase model. This model consists of 30 species and 54 parameters. For this model, we predict various parameter values from the characterization of the integrase circuit in cell-free extract. We use the identified and validated parameters for the cell-free extract resources, integrase expression, and integrase action in the excisionase model. The integrase action parameters in this model are context-dependent, so, we allow for these to be updated as we validate using the integrase-excisionase cell-free data. However, we keep the cell-free resource parameters constant as they model the total resources provided by the extract.

We analyze the sensitivities of each parameter in this model to the output and run simulations under various conditions to predict the excisionase action *a* priori. This guides our design choices of choosing the excisionase plasmid levels relative to the integrase plasmid levels. As we vary integrase plasmid initial conditions from 0 to 2 nM, we observe that varying the excisionase initial conditions from 0 to 1 nM gives accurate control of output protein desired levels of expression. This is the design choice we made for the cell-free experiments with both integrase, excisionase, and reporter plasmids (resulting *in vitro* data shown in Figure 9). These forward design choices driven by mathematical models were made possible due to the characterization of a smaller part of this circuit (the integrase-reporter circuit), the detailed models, their sensitivity analyses, and with preliminary excisionase-reporter cell-free experiments.

To validate the mathematical models by identifying parameter values from the experimental data, we need to reduce the dimension of the parameter space of the detailed model. Although the detailed model captures a range of behavior, such as resource loading and competition, it is infeasible for parameter identification due to the unidentifiability of its large parameter space. So, we use AutoReduce to reduce the models in multiple steps.

This model reduction process is shown in Figure 8 which starts at automatically deriving and applying the conservation laws, then proceeds to QSSA, and finally, context abstraction with BioCRNpyler to derive a minimal ODE model (M-8 in Figure 8). This minimal model is suitable for parameter identification as it contains only 5 states and 17 parameters out of which 4 parameters are most sensitive to YFP expression (see Figure 9B). Using the identified parameters, we are able to characterize and predict the excisionase activity in reversing the integrase action, their relative strengths, and the ratios required for accurate protein level prediction. The experimental data and the parameter identification steps for excisionase characterization are shown in Figure 9.

## Conclusion

We present a new *in silico* pipeline to assist the design-build-test-learn process in synthetic and systems biology. This pipeline consists of user-friendly, open-source, Python software packages that have been developed with community input and support standardized models written in SBML. We build on three Python packages to formulate the pipeline presented in this paper, which starts by building detailed reaction network models for biological circuits, reducing the detailed models to simpler ODEs, and seamlessly connecting these models to experimental data through Bayesian inference. All software packages demonstrated in this paper are Python-based (for easy integration with other tools and pipelines) and have an active discussion and issue support forums through Github.

We demonstrate the application of this pipeline with a novel circuit design in a cell-free protein synthesis system. We build a three-plasmid system consisting of DNA recombination enzymes, integrases, and excisionases. With the help of this pipeline, we characterize the expression and action of integrases and their reverse directionality factors, excisionases. This characterization involves detailed reaction network models, ODE models under various assumptions with a clear indication of when each model is valid, and posterior parameter distributions for model parameters that fit the experimental data. We quantify the expression strengths of integrase, excisionase, and the reporter while also quantifying their relative effects on the output fluorescence measurement. Particularly, we show control of output protein expression at various levels with experimental data that is backed by mathematical models. We postulate that predictable DNA recombination-based control of protein expression adds an alternative design choice for biological circuit designs. This characterization of circuit parts in the cell-free system with mathematical models would allow for such complex circuit designs that use integrases and excisionases for precise control of protein expression. However, further characterization and experimentation, especially concerning the loading effects, are required to achieve that goal. The computational pipeline that we have developed is general enough to assist these future research directions.

There are two noteworthy limitations of the work presented in this paper, both of which are also open research problems in systems and synthetic biology:

1. Scalable biological circuit design: Although the computational pipeline presented in this paper applies to larger biological circuits as well, it is only demonstrated for circuits with 3-4 components. Various challenges limit our ability to quantify and model larger circuits. From the biological standpoint, it is still unclear how the context changes and affects the performance when more components are added to a system. On the other hand, parameter identification for higher dimensional problems is computationally inefficient, which presents another bottleneck in validating larger circuit models.
2. We show that reduced models can be systematically obtained from detailed models for computationally feasible parameter inference. But, in some cases, we observe that the reduced models are unable to fit the experimental data (for example, the plateauing of fluorescence as the cell-free extract runs out of resources). We expect to further build on the modeling tools presented in this paper and integrate these with other tools and pipelines to address some of these issues.

## Methods

### Cell-free Experiments

Experimental characterization of both integrase and excisionase activity was done in a cell-free extract – an *in vitro*, cell-free protein expression system. As depicted in Figure 2, the cell-free reactions created are composed of S30 *E. coli* cell extract, energy buffer, and DNA of our designed genetic circuit.^31^ The cell-free extract and energy buffer were prepared following the protocol in Sun et al.^32^ Plasmid DNA was added to a cell-free master mix of cell extract and energy buffer to create each reaction. When analyzing the integrase activity, we used automated acoustic handling (Labcyte Echo 525) to load reagents into a 96-well plate and vary the level of the integrase and reporter plasmids from 0 to 1 nM over 5 concentrations: 0 nM, 0.25 nM, 0.50 nM, 0.75 nM, and 1 nM. Each reaction contained one possible combination of the integrase plasmid and reporter plasmid, creating a total of 25 reactions. A total of 50 reactions were generated to achieve duplicates for all possible reactions. After all of the reactions had been created, they were incubated at 29°C while measuring the CFP (Ex: 440; Em: 480) and YFP (Ex: 503; Em: 540) fluorescence every 5 minutes in a Biotek Plate Reader.

When analyzing the excisionase activity, we varied the level of the integrase plasmid and excisionase plasmid from 0 to 2 nM and from 0 to 1 nM, respectively, while keeping the concentration of the reporter plasmid constant at 1 nM for each reaction. Again, using automated acoustic handling (Labcyte Echo 525), we varied the concentration of each plasmid independently over 5 concentrations. The integrase plasmid was added to each reaction at 0 nM, 0.5 nM, 1.0 nM, 1.5 nM, or 2 nM. Additionally, the excisionase plasmid was added at 0 nM, 0.25 nM, 0.5 nM, 0.75 nM, or 1 nM. Each reaction contained one possible combination of the integrase plasmid and reporter plasmid, creating a total of 25 reactions. A total of 50 reactions were generated to achieve duplicates for all possible reactions. After all of the reactions had been created, they were incubated at 29°C while measuring the CFP (Ex: 440; Em: 480), RFP (Ex: 564; Em: 594), and YFP (Ex: 503; Em: 540) fluorescence every 5 minutes in a Biotek Plate Reader.

### Model Reduction Methods

We used a variety of model reduction techniques to derive reduced models in this paper. We chained these methods in an automated workflow by developing the Python model reduction software, AutoReduce.^33^ First, all CRN models with mass-action kinetics admit a set of conservation laws. The underlying theory and the derivation of conservation laws is a well-studied topic in CRN theory.^34^ We implemented Python methods in AutoReduce to search the conservation laws in a given model (not necessarily a mass-action CRN), computation of reduced models with conservation law substitutions, and symbolic manipulation of the model as well as numerical computations for symbolic models. For CRNs with mass-action kinetics, we find that the derived conservation laws with AutoReduce are mass conservation laws, such as the total RNA polymerase being conserved as the sum of free polymerase, and all species in the model with a bound polymerase. Similarly, we have conservation laws for total DNA, total ribosomes, total endonucleases, and other resources.

After applying conservation laws, we used quasi-steady state approximation (QSSA) to further reduce the models. The built-in automated QSSA tools in AutoReduce were used for this purpose. By applying QSSA iteratively to reduced models, we obtain further reduced models. The details of which species were assumed to be at QSS are shown in Figures 4 and 8. For both full models (the integrase action and integrase-excisionase action), we obtain a set of reduced models, each with different assumptions in the derivation of the reduced model. These are shown as flowcharts in Figures 4 and 8. Some reduced models were amenable to even further reduction in dimensionality by introducing species abundance assumptions. For example, on assuming abundance of RNA polymerase (*P*_tot_), we may write the terms like *P*_tot_ − *C* as approximately equal to *P*_tot_ where *C* is an intermediate low concentration complex species. Such assumptions were only true under certain parameter conditions and for certain reduced models, hence, did not always lead to correct reduced models (see the section on performance metrics in Methods for more on how we validate a reduced-order model). Nevertheless, with the Python tools in AutoReduce and our further additions to it, exploration of the space of models is quick. Finally, we extended AutoReduce to develop wrappers for easy import and export of SBML files so that the tool can be seamlessly connected to existing pipelines.

### Context Abstraction with BioCRNpyler

BioCRNpyler allows the modeling of biological systems in different contexts and modeling details with the use of built-in `Mixtures` (objects that model the context-dependent and global effects) and `Mechanisms` (objects that capture the modeling detail for a process). In this paper, we utilize this key functionality of BioCRNpyler to explore the design space and possible hypothesis for integrase and excisionase-based DNA recombination. We developed two integrase action mechanisms — a simple mechanism that models the flipping of attP-attB promoter sequence to attL-attR in one step, and a detailed mechanism that models the same process but with binding events involving integrase binding to the different DNA sites. For excisionase action, the exact mechanism and binding steps are unknown, so we model all possible steps and explore the hypothesis with our iterative pipeline and the experimental data. We model the excisionase binding to DNA already bound with integrase on the attP-attB or the attL-attR sites as the first hypothesis of the excisionase mechanism. Once excisionase binds at the attL-attR site already bound with integrase, it flips the sequence to attP-attB. Another potential mechanism by which excisionase may reverse the directionality of the promoter is by binding to the integrase first and forming an integrase-excisionase complex. This complex, when bound with the DNA at the attL-attR site, flips the promoter to attP-attB. Further, the sequestration mechanism for excisionase is also modeled by including the reactions in which the excisionase binds to the attP-attB site, hence, preventing integrase to act on it. We add all of these mechanisms to the library of BioCRNpyler mechanisms so that they can be used to build CRN models in different mixtures. We use the `TxTlExtract` mixture in BioCRNpyler along with the mechanisms described above to build detailed CRN models. In the iterations of the pipeline, as described in Figure 4 and 8, we switch mechanisms `ON` or `OFF` to abstract the details of the model and switch `Mixtures` to build simpler or detailed models. In conclusion, the context abstraction with BioCRNpyler is achieved by building mechanisms and mixtures and simply choosing the mixture to use for context and the mechanisms for the modeling detail.

### Reduced Model Performance Metrics

Each reduced model obtained in this paper was tested for its performance against the full model by computing different metrics. We used the metrics developed in Pandey et al.^15^ for this purpose. First, we computed the normed error metric, *ζ*, for each reduced model:

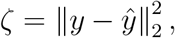

where *y* and 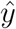 are the outputs of the full and the reduced model respectively. In all cases analyzed in this paper, we derived reduced models where the outputs of the full model were never collapsed since these are the signals which are measured. For most cases, the error performance metric sufficed in rejecting the reduced models, however, two other metrics that were shown to be useful in deciding among equally good error-performance models are:

1. Error sensitivity: It was shown that when normed error performance is of the same order for two or more reduced models, we can compute the sensitivity of the error with respect to each parameter in the model to choose a reduced model whose error is least sensitive to parameter perturbations.
2. Input-output map: For each reduced model, we computed an input-output system gain using linear systems theory^35^ by first linearizing the model around a point of interest. This input-output gain was then compared with the gain of the full model to ensure the fidelity of the reduced models.

Using these error metrics, we decided which reduced model to choose as shown in Figures 4 and 8. The computation of the metrics for all reduced models shown in this paper is available with the associated Python code on Github.^36^

### Data Analysis and Parameter Inference

Standard Python libraries (NumPy^37^ and Matplotlib^38^) were used for the analysis and plotting of experimental data. Three main analysis and optimization tools were used in this paper:

#### Numerical Simulations

The Cython-based tool Bioscrape^16^ provides access to fast simulators suited to simulate CRN models even under stiff conditions that are commonly observed in mass-action ODE simulations. Although Bioscrape provides both deterministic and stochastic simulation tools we used only deterministic simulations for the analyses in this paper. The Python wrappers available in Bioscrape were used to import SBML files from BioCRNpyler and AutoReduce to run simulations. All simulations shown in this paper were performed using Bioscrape.

#### Sensitivity Analysis

We extended the suite of analysis tools in Bioscrape by developing local sensitivity analysis methods. The local sensitivity coefficients are computed at each time point as

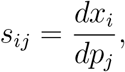

where *x_i_* is the i-th state and *p_j_* is the j-th parameter. The sensitivity coefficients are arranged in a tensor of size length of timepoints × number of parameters × number of states. The computation of sensitivity coefficients is done by using the direct method.^39^ Various options to control the accuracy, normalization, and integration methods are available to the user. Full documentation of the sensitivity analysis was added to the Bioscrape documentation on Github.^40^ In Figures 3 and 7, we used the sensitivity analysis method to assess which parameters were most effective for each model. In Figures 5 and 9, we used the sensitivity analysis to find identifiable parameters from the data and guide a by-parts parameter inference process.

**Figure 7:**
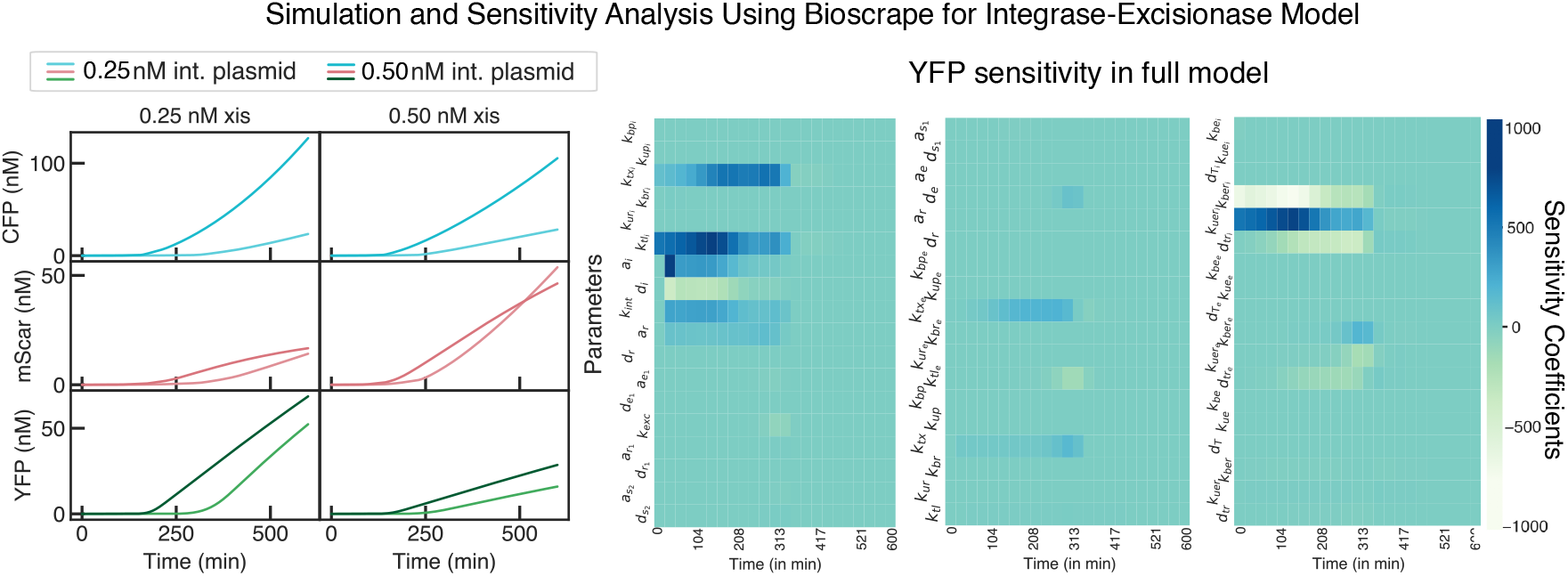
Analysis for excisionase expression and activity in cell-free extract using Bioscrape. We show the simulation for the CRN model. In this model, we use the identified integrase parameters to predict the excisionase activity. We observe that as more excisionase is expressed, YFP expression falls down. The sensitivity of each parameter in the CRN model with time is shown in the sensitivity analysis heatmaps.

**Figure 8:**
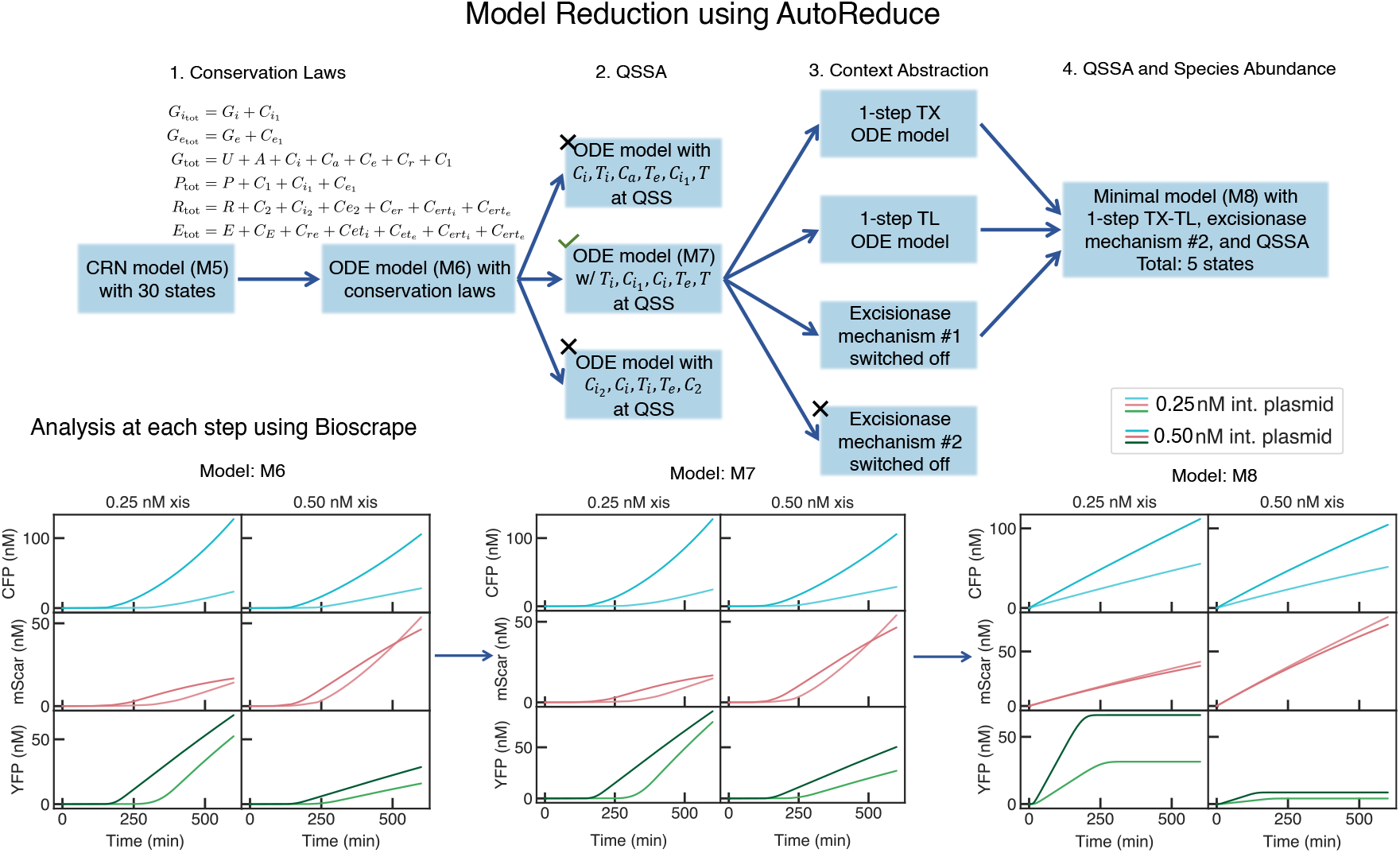
Mathematical models for excisionase expression and activity in cell-free extract obtained using AutoReduce and BioCRNpyler. Using AutoReduce, we find the conservation laws in the CRN model given by BioCRNpyler. After applying conservation laws, we further reduce the dimensionality of the integrase-excisionase system by assuming species at a quasi-steady state (QSS). Finally, we abstract the context of resource details modeled using BioCRNpyler and reduce these models further using AutoReduce to obtain a minimal model (M8). The chosen reduced models are marked with a tick while a cross indicates discarded reduced model. The simulations for three reduced models are shown — M6, M7, and M8. The computational run time to derive these reduced models are similar to the integrase reduced models, varying from a few seconds to a few minutes.

**Figure 9:**
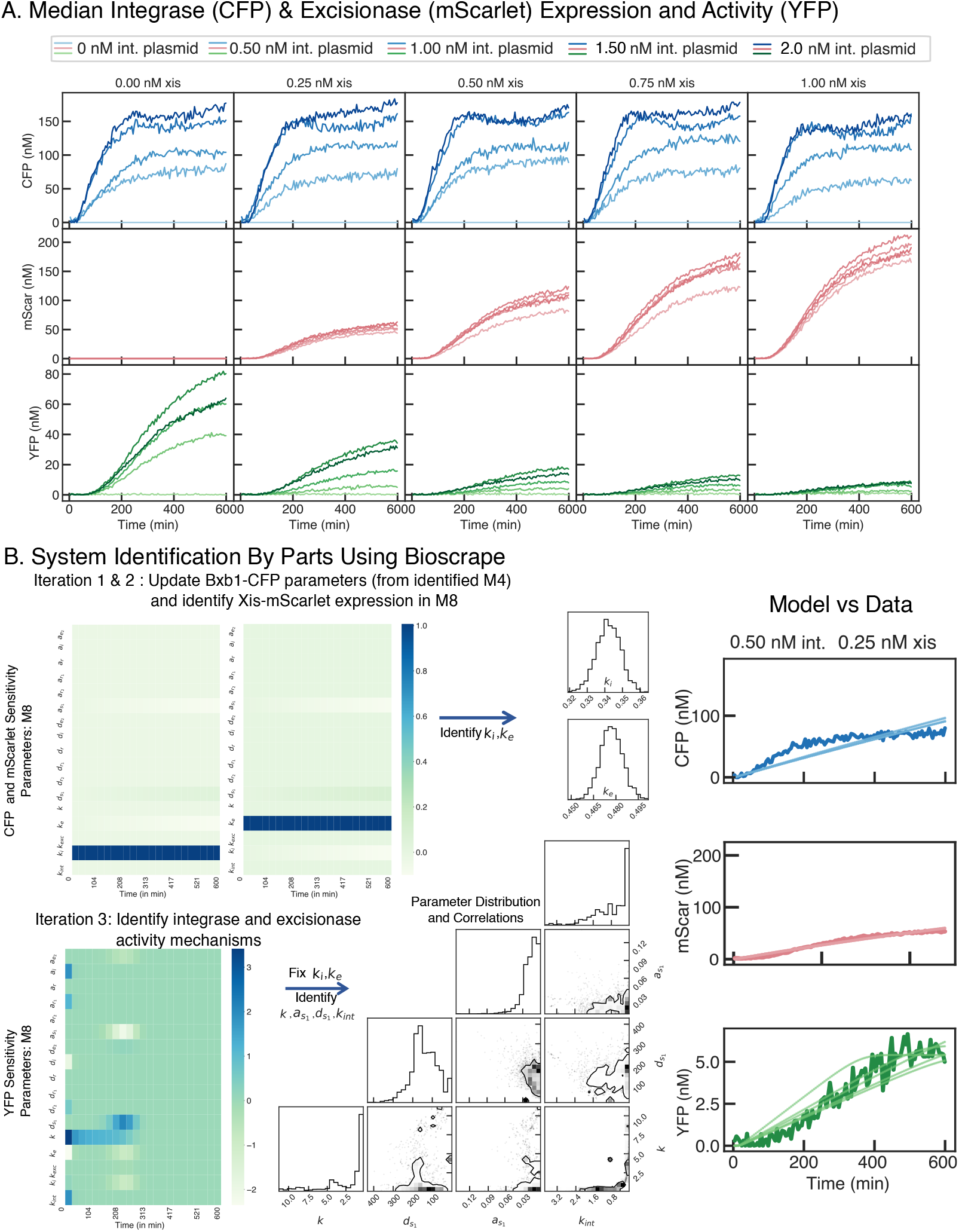
Experimental data and system identification by parts of integrase and excisionase expression and activity in cell-free extracts. (A) Median background-subtracted fluorescence data for the integrase-excisionase circuit in cell-free extract. We observe that as more integrase is added, more YFP is expressed until the maximum possible expression is reached at 1.5nM integrase. With higher excisionase levels, we observe a decreasing gradient of YFP levels. (B) To identify the model parameters, we set the previously identified integrase mechanism parameters and update those accordingly to account for various loading effects. In the second iteration, we infer the parameters for the mScarlet expression. Finally, we identify the sensitive set of parameters for YFP. In the right panels, we show the identified parameter distributions and the data plotted alongside the model simulations. For more details on parameter inference, see the supplementary note on parameter inference.

#### Bayesian Inference Using MCMC

We used time-series fluorescence reporter measurements to validate the mathematical models. Towards that end, we optimized the parameters in the model using Bayesian inference. We find probability distributions for parameters given the experimental data (posterior distributions) and then sample the parameter values from the posterior to simulate the model to plot alongside the data. Bayes’ rule is the underlying principle for this approach:

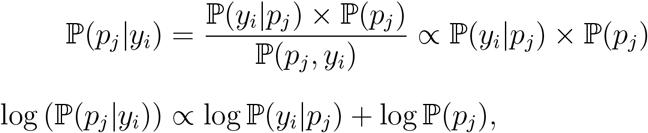

where *p_j_* consists of the parameters for which we want the probability distribution given the data instance(s) *y_i_*. The probability distribution ℙ(*y_i_*|*p_j_*) is called the model likelihood since it is proportional to the likelihood of seeing the data *y_i_* given that the parameters take the value *p_j_*. We implemented multiple ways in which this likelihood can be computed by simulating the model at the given parameter values *p_j_* and comparing the model against the data. For example, the likelihood may be computed with Bioscrape as the normed difference between the model outputs and experimental measurements over time or it may be computed as the maximum error between the simulation and the data. We implemented a total of 6 methods to compute the model likelihood. The Bioscrape documentation describes these in detail. For the parameter inference in this paper, we used the deterministic trajectory likelihood which computes the normed error between the model and the data for all time in the data trajectories. The probability distribution ℙ(*p_j_*) is the prior probability distribution that gives each parameter value a probability of being the true value from the prior information about the parameters. We implemented multiple prior probability distributions in Bioscrape including, Gaussian, uniform, log-normal, log-uniform, beta, and more. A custom function may also be used as a prior probability distribution for a parameter. Bioscrape documentation describes their usage in detail. Since all models used in this paper are models that describe biophysical processes, the parameters have mechanistic meanings, hence, priors were used to constrain the parameter inference according to the biological prior knowledge about each parameter. To compute each parameter sample, we use the Python emcee^17^ package that implements a Markov Chain Monte Carlo (MCMC) sampler. This MCMC sampler proposes the next parameter sample by assessing how the previous sample performed. For more details on the sampling algorithm, the reader is referred to the emcee documentation.^17^ In conclusion, we developed Bioscrape inference as a black-box wrapper that imports experimental data and an SBML model to be used for parameter inference with emcee. Code for all data analysis, parameter inference, and related documentation are available on Github.^36^

## Acknowledgement

The authors thank the many users of BioCRNpyler, AutoReduce, and Bioscrape who have provided invaluable feedback on the software tools demonstrated in this paper. In particular, the authors acknowledge Victoria Hsiao for providing the integrase plasmid used in this paper, Anandh Swaminathan for working on a preliminary version of the integrase model, Andrey Shur and Zoltan Tuza for their feedback on the analysis, and Zoila Jurado and Alex Johnson for their insightful comments on the manuscript.

A.P. is currently supported by the AFOSR MURI grant FA9550-22-1-0316 and was previously funded in part by the NSF grant CBET-1903477 and DARPA grant HR0011-17-2-0008. W.P. was partly supported by the NSF grant MCB-2152267 and Army Research Office grant W911NF-19-D-0001. The content of the information does not necessarily reflect the position or the policy of the Government, and no official endorsement should be inferred.

## Supporting Information Available

### Raw Data and Concentration Calibration

The raw fluorescence data for the integrase and reporter circuit shown in Figure 2 is shown in Figure 10. For the excisionase expression (mScarlet fluorescence) and its action on the reporter, the raw fluorescence data for CFP (integrase expression), mScarlet (excisionase expression), and YFP (reporter) is shown in Figure 11. We calibrated our plate reader reading to obtain absolute fluorescence measurements for the cell-free experiments. The calibration factors are presented in Table 1 where arbitrary units [AU] = Slope · [uM] + Offset.

**Figure 10:**
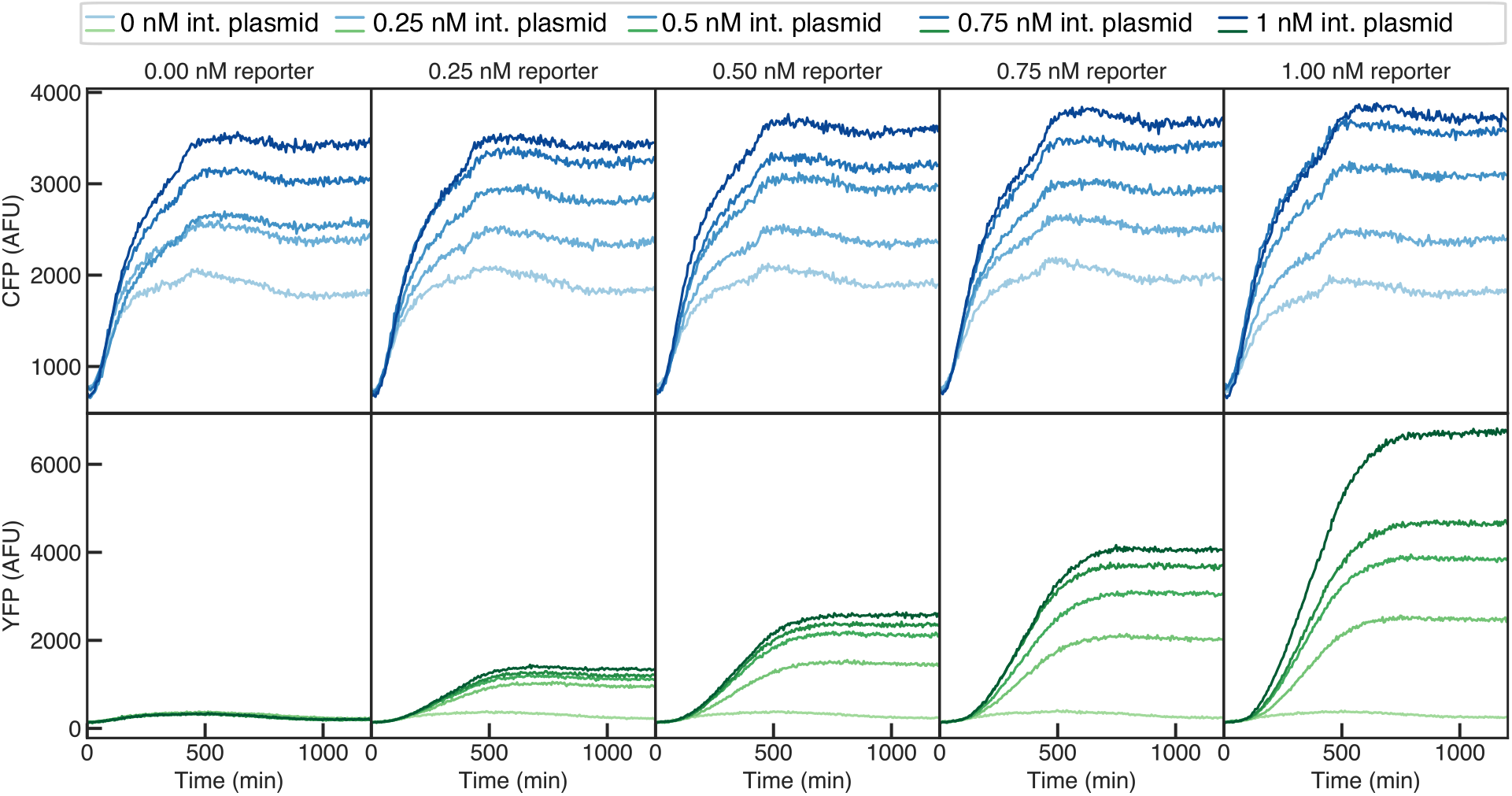
Cell-free fluorescence data for the integrase and reporter expression circuit. The raw measurements are in arbitrary units. We process these measurements by subtracting the background and calibrating the fluorescence to concentration units (details of the cell-free experiments are given in the Methods section).

**Figure 11:**
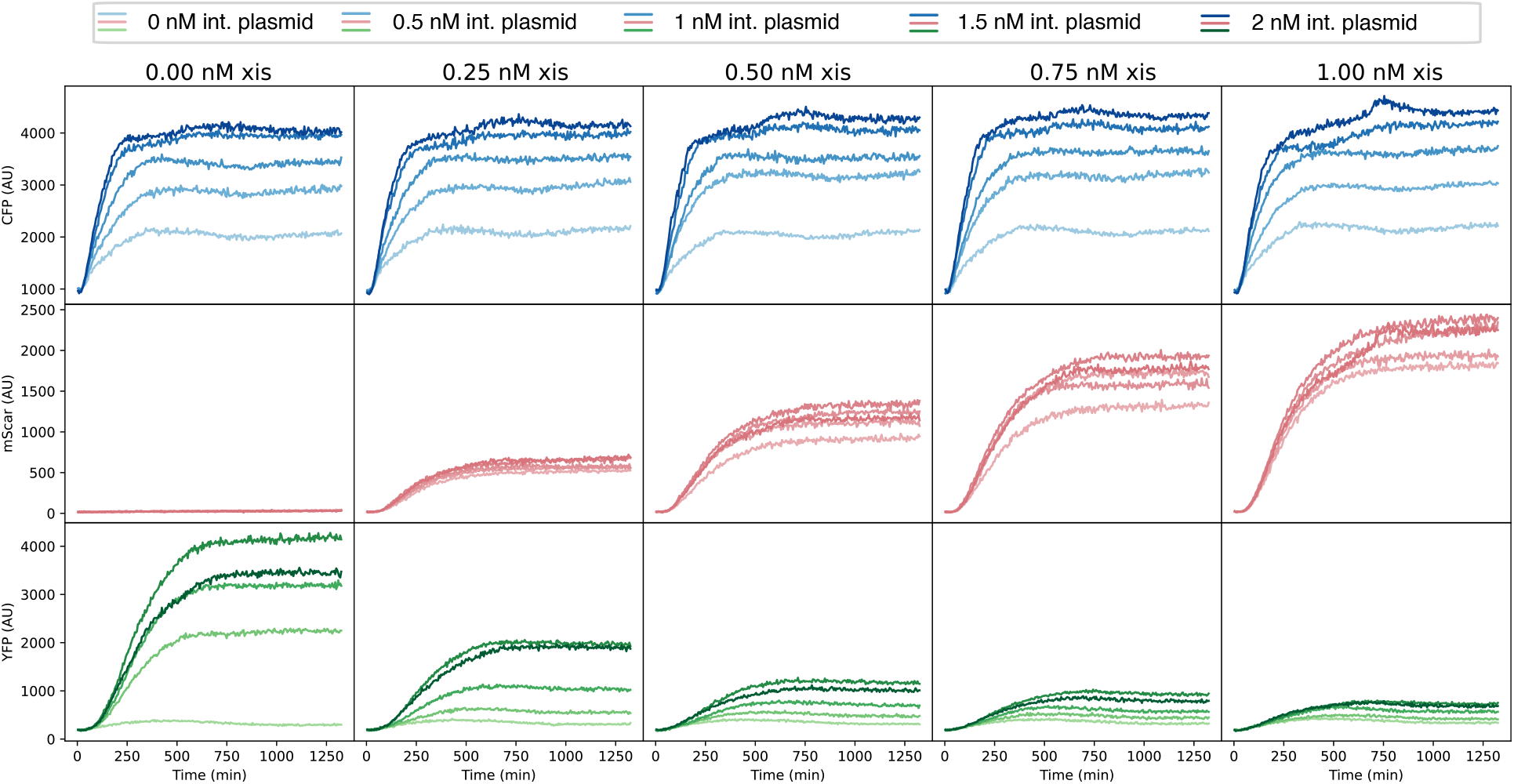
Cell-free fluorescence data for the integrase, excisionase, and reporter expression circuit. The raw measurements are in arbitrary units. We process these measurements by subtracting the background and calibrating the fluorescence to concentration units (details of the cell-free experiments are given in the Methods section).

**Table 1:** Calibration factors for cell-free experiments

| Measurement | Slope | Offset |
| --- | --- | --- |
| CFP | 12796.23 | -6248.64 |
| mScarlet | 9886.58 | 166.59 |
| YFP | 44451.55 | 294 |

**Table 2:**
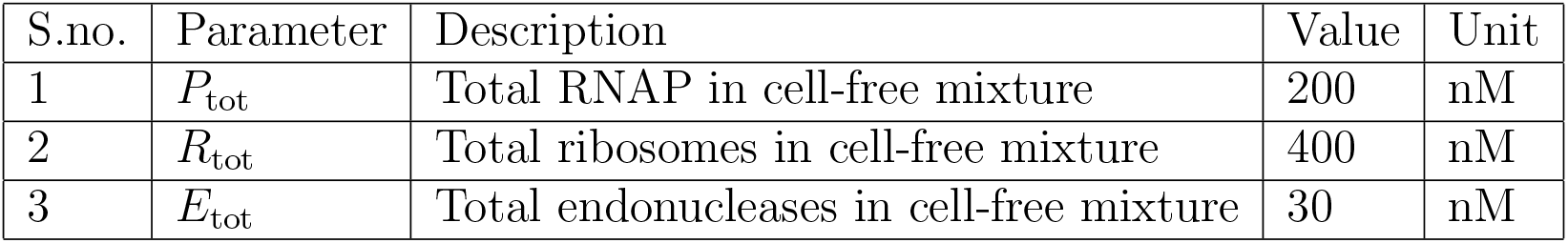
Cell-free global resources

**Table 3:** Key integrase expression and action parameters in cell-free extract

| S.no. | Parameter | Description | Value | Unit |
| --- | --- | --- | --- | --- |
| 1 | $k_{tx_i}$ | Transcription rate of integrase expressing mRNA | 0.25 | hour<sup>-1</sup> |
| 2 | $k_{tl_i}$ | Translation rate of integrase-CFP expression | 0.013 | hour<sup>-1</sup> |
| 3 | $n$ | Cooperativity of integrase action | 3 | – |
| 4 | $k_{int}$ | Integrase flipping rate | 0.1 | hour<sup>-1</sup> |
| 5 | $k_{tx}$ | Transcription rate of the reporter-mRNA | 0.3 | hour<sup>-1</sup> |
| 6 | $k_{tl}$ | Translation rate of the reporter | 0.03 | hour<sup>-1</sup> |

Note: Only 6 out of 29 parameters are given in the table.

**Table 4:** Key integrase-excisionase expression and action parameters in cell-free extract

| S.no. | Parameter | Description | Value | Unit |
| --- | --- | --- | --- | --- |
| 1 | $k_{tx_i}$ | Transcription rate of integrase expressing mRNA | 0.29 | hour<sup>-1</sup> |
| 2 | $k_{tl_i}$ | Translation rate of integrase-CFP expression | 0.008 | hour<sup>-1</sup> |
| 3 | $n_i$ | Cooperativity of integrase action | 3 | – |
| 4 | $k_{int}$ | Integrase flipping rate | 0.1 | hour<sup>-1</sup> |
| 5 | $k_{tx_e}$ | Transcription rate of excisionase expressing mRNA | 0.292 | hour<sup>-1</sup> |
| 6 | $k_{tl_e}$ | Translation rate of excisionase-mScarlet expression | 0.011 | hour<sup>-1</sup> |
| 7 | $n_e$ | Cooperativity of excisionase action | 4 | – |
| 8 | $k_{exc}$ | Excision rate | 0.7 | hour<sup>-1</sup> |
| 9 | $k_{tx}$ | Transcription rate of the reporter-mRNA | 0.4 | hour<sup>-1</sup> |
| 10 | $k_{tl}$ | Translation rate of the reporter | 0.08 | hour<sup>-1</sup> |

Note: Only 10 out of 54 parameters are shown. Note that some integrase parameters are updated in the parameter identification process as the context changes in this more complex circuit

### Model Species

In the integrase model (shown in Figure 2), each variable represents the concentration of the model species. *G_i_* is the integrase plasmid, *P* is RNA polymerase (RNAP) and *C_i_*1 is the binding complex between RNAP and the gene. *T_i_* is the mRNA transcript that codes for the Bxb1-CFP (*I*) protein. The binding complex between mRNA and ribosome, *R*, is denoted as 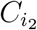. The reporter plasmid is called *U* in its “inactive” state (that is, when the promoter is in the reverse direction) and *A* in its “active” state. RNAP binds to the reporter gene to form the complex *C*_1_ to transcribe the reporter mRNA *T*, which then binds to the ribosome (*R*) to form *C*_2_. The YFP concentration is denoted as *Y*. For the integrase activity, we use *n_i_* for the cooperativity of integrase binding to the inactive gene, *U*, to form the complex, *C_i_*. The integrase flipping reaction forms the active complex, *C_a_*, which then reversibly forms the active reporter plasmid, *A*. We model the degradation of mRNA and its complexes by the endonucleases in cell-free extracts. The endonuclease concentration is denoted by *E*, which forms complexes 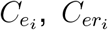, *C_e_*, and *C_er_* when it binds to the integrase mRNA transcript, the ribosome-bound integrase mRNA transcript, the reporter transcript, and the ribosome-bound reporter mRNA transcript respectively.

In the integrase-excisionase model (shown in Figure 6), we have the transcription and translation of the Xis-mScar gene (*G_e_*) that expresses the fused excisionase-mScarlet protein (*E*_0_). The corresponding transcript is denoted as *T_e_*, the RNAP-gene complex as 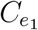, and the transcript-ribosome binding complex as 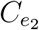. For the excisionase action, we introduce two mechanisms. In the first mechanism, the excisionase, with a cooperativity of *n_e_*, binds to the integrase-bound “active” reporter plasmid, *C_a_*, with attL-attR sites to form the complex *C_e_*. The excisionase flips the promoter direction in this complex to form *C_r_*, which is the “inactive” complex with both integrase and excisionase bound at the attB-attP site. Then, this complex reversibly unbinds to give out free integrase, excisionase, and the “inactive” reporter plasmid, *U*. In the second excisionase mechanism, the excisionase binds to the integrase forming the complex *C_ie_*. The excisionase may also bind to the integrase-bound DNA, *C_i_*, forming *C_r_*, to sequester the integrase from flipping *C_i_* to *C_a_*. Finally, the integrase-excisionase complex, *C_ie_*, binds to both the “inactive” and the “active” DNA (*U* and *A*) to form complexes *C_r_* and *C_e_* respectively. Similar to the endonuclease-mediated degradation of integrase and reporter mRNA and their complexes, the excisionase mRNA and its complexes are also degraded. The endonuclease binds to the excisionase mRNA to form 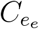 and it binds to the ribosome-bound excisionase mRNA, 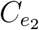, to form the complex 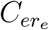.

### Parameter Values

All parameter values and simulation conditions for all models in this paper are available on Github.^36^ To summarize some of these findings, we provide key parameter values for the detailed and minimal models as tables below. Note that we identified the posterior distributions (and hence, the uncertainties) for the parameters in the minimal models. The corner plots in Figures 5 and 9 show these uncertainties in each identified parameters. The reader is referred to the source code^36^ for more details on statistical convergence, prior distributions, initial guesses, and posterior distributions with uncertainties.

### Parameter Inference and Unidentifiability

#### MCMC Sampler for Inference of Integrase-Reporter Circuit

In this section, we show the parameter inference results in detail for the integrase model and the integrase-excisionase model. The parameter inference uses experimental data from cell-free experiments. The same cell-free mixture (extract and buffer) is used for all experiments shown in this paper to ensure that the context-dependent parameters remain constant throughout.

For the integrase expression and action (model shown in Figure 2), the MCMC chains for the identified parameters are shown in Figure 12. As discussed in the main text, we use system identification by parts to identify parameters sequentially using the sensitivities of the measurements to different parameters. The first step of parameter identification uses CFP measurement to identify *k_i_* as suggested by sensitivity analysis (see Figure 5B). We run the MCMC sampler for 1000 steps, 10 walkers, and a Gaussian prior on *k_i_* with a mean value of 0.23 and a standard deviation equal to 1. To identify the integrase action and reporter expression parameters, *k_int_*, *n*, *K_I_*, and *k*, we use YFP measurement as suggested by the sensitivity analysis. For this MCMC sampler, we use 40 walkers for 10000 steps and Gaussian priors on all parameters. The mean values used in the priors for *k_int_*, *n*, *K_I_*, and *k* are 0.05, 2, 3330, and 0.0001 while the standard deviations used are 10, 2, 1000, and 0.1 respectively. The total runtime for this parameter inference was 50 minutes on a personal computer with an Intel i7-6700 processor and 16GB of RAM.

**Figure 12:**
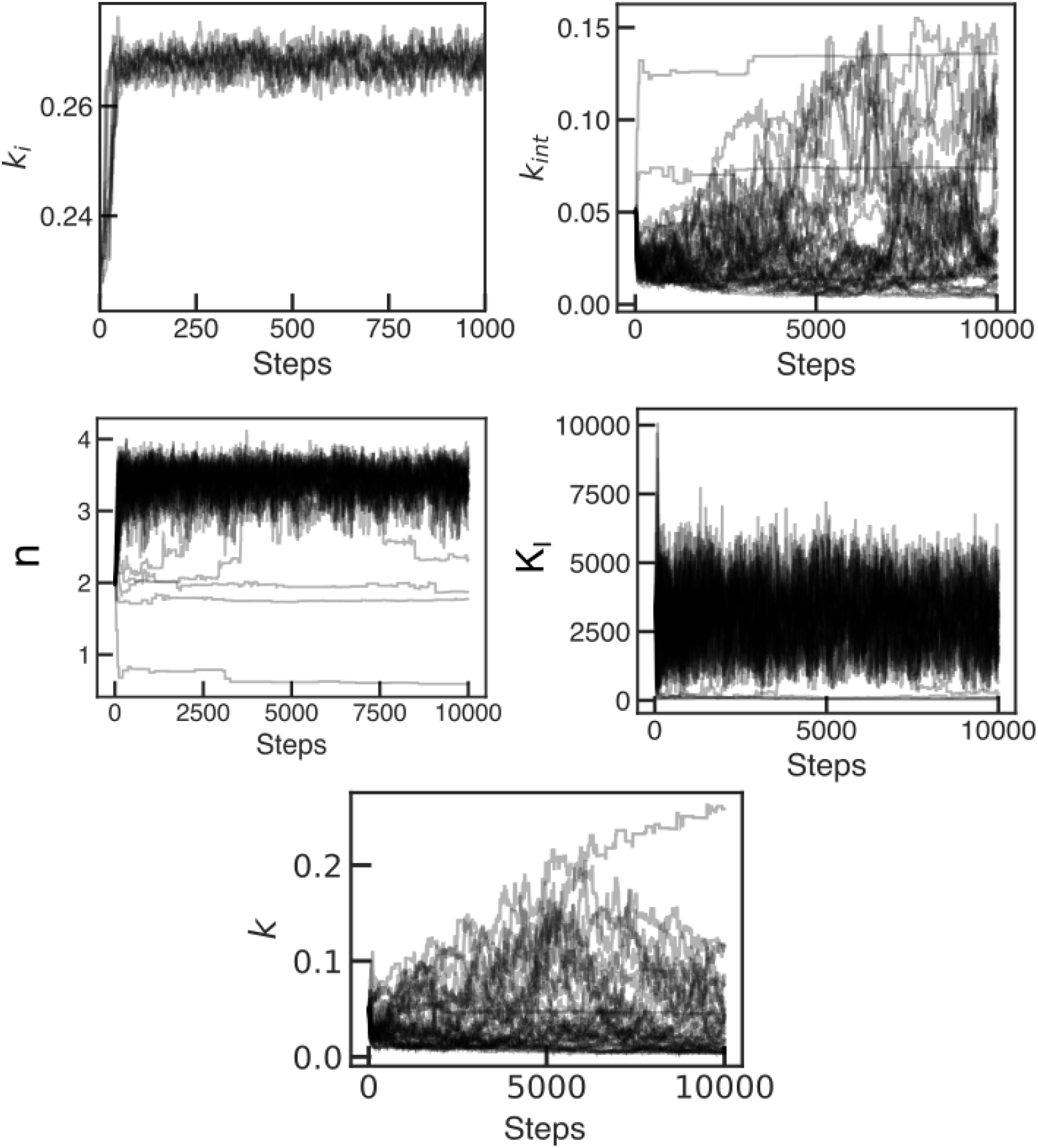
MCMC chains for identification of integrase model parameters given the integrase-reporter cell-free experiments.

#### Identified Integrase Model and Data

We sample from posterior parameter distributions and run model simulations. We plot the simulations of the identified models together with the experimental data. These results are shown for both the CFP and YFP measurements in Figures 13 and 14 respectively.

**Figure 13:**
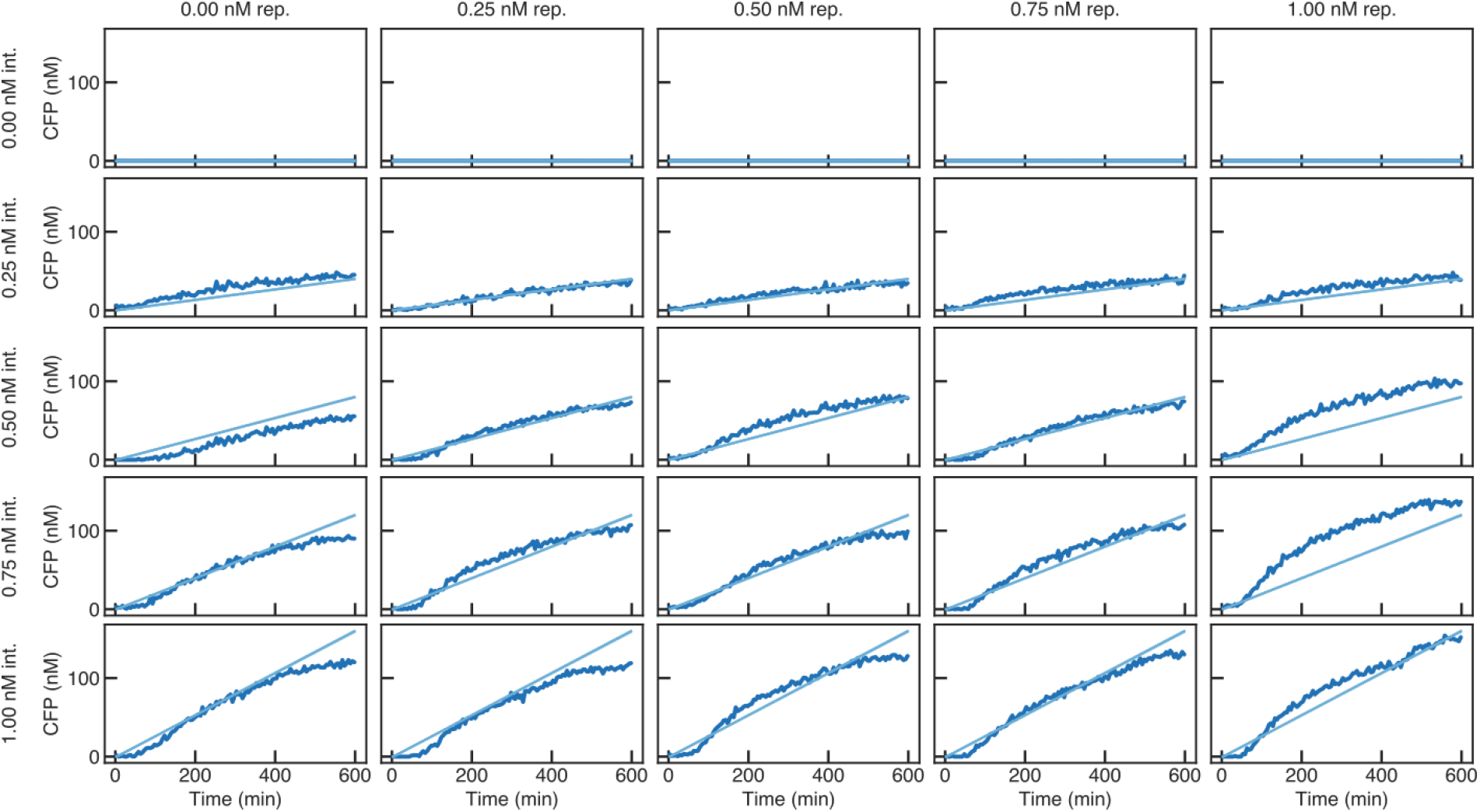
Model simulations with posterior parameter values plotted alongside CFP measurement. Observe that the model fits the data well for most conditions, however, for higher reporter plasmid concentration the fit worsens. See the discussion on loading effects in Material and Methods for more information on this observation.

**Figure 14:**
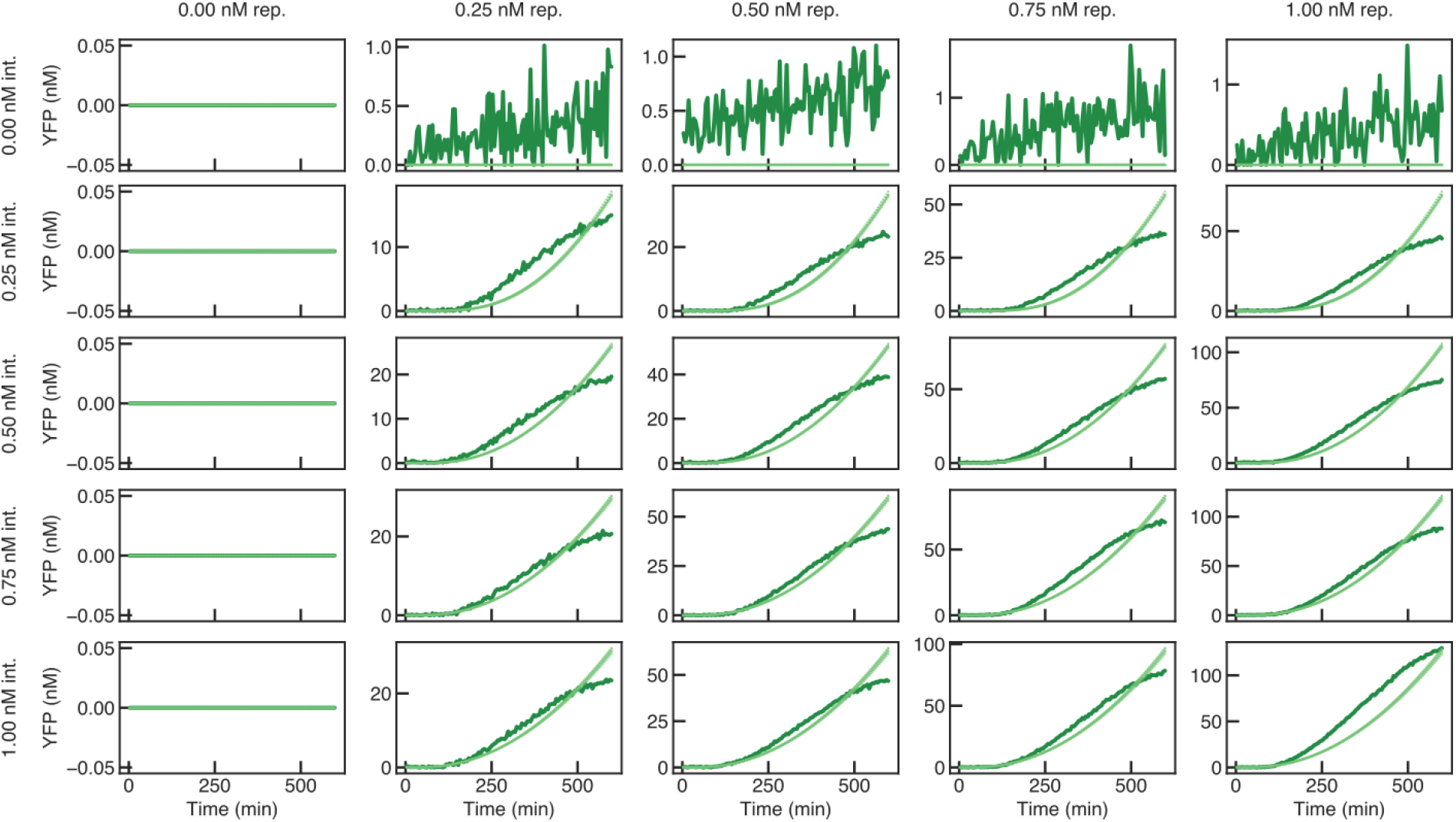
Model simulations with posterior parameter values plotted alongside YFP measurement. Observe that the minimal model (derived automatically from a detailed CRN model) does not fit the dynamics as cell-free extract stops protein expression. This is a result of context abstraction steps in obtaining this minimal model. Since the effects of resource usage are not modeled in this minimal model, the YFP expression in the model does not saturate as quickly as the observed data.

#### MCMC Sampler for Inference of Integrase-Excisionase Circuit

Here we describe the MCMC sampler used to infer the parameters of the integrase-excisionase and reporter circuit (model shown in Figure 6). The MCMC chains for the identified parameters are shown in Figure 15. First, we use the CFP measurement to re-estimate *k_i_* in this updated context and then use the mScarlet measurement to infer *k_e_*. See sensitivity analyses of the different outputs (CFP, mScarlet, and YFP) to model parameters in Figure 7. We run the MCMC sampler for 1000 steps, 10 walkers, and Gaussian priors on *k_i_* and *k_e_* with mean values set at 0.26 (previously identified maximum likelihood value) for *k_i_* and 0.3 for *k_e_*. Both priors are used with a standard deviation equal to 5. Further, to identify the integrase and excisionase action, and reporter expression parameters, *k_int_*, *d_i_*, 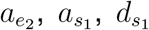, and *k*, we use YFP measurements as suggested by the sensitivity analysis. For this MCMC sampler, we use 20 walkers for 20000 steps and Gaussian priors on all parameters. The mean values used in the priors for *k_int_*, *d_i_*, 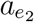, *k_e_*, 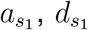, and *k* are 0.2, 500, 0.02, *k_e_* (previously identified), 0.1, 100, and 2 while the standard deviations used are 5, 1000, 5, 0.1 ∗ *k_e_*, 5, 100, and 10 respectively. The total runtime for this parameter inference was 24 hours on a personal computer with an Intel i7-6700 processor and 16GB of RAM.

**Figure 15:**
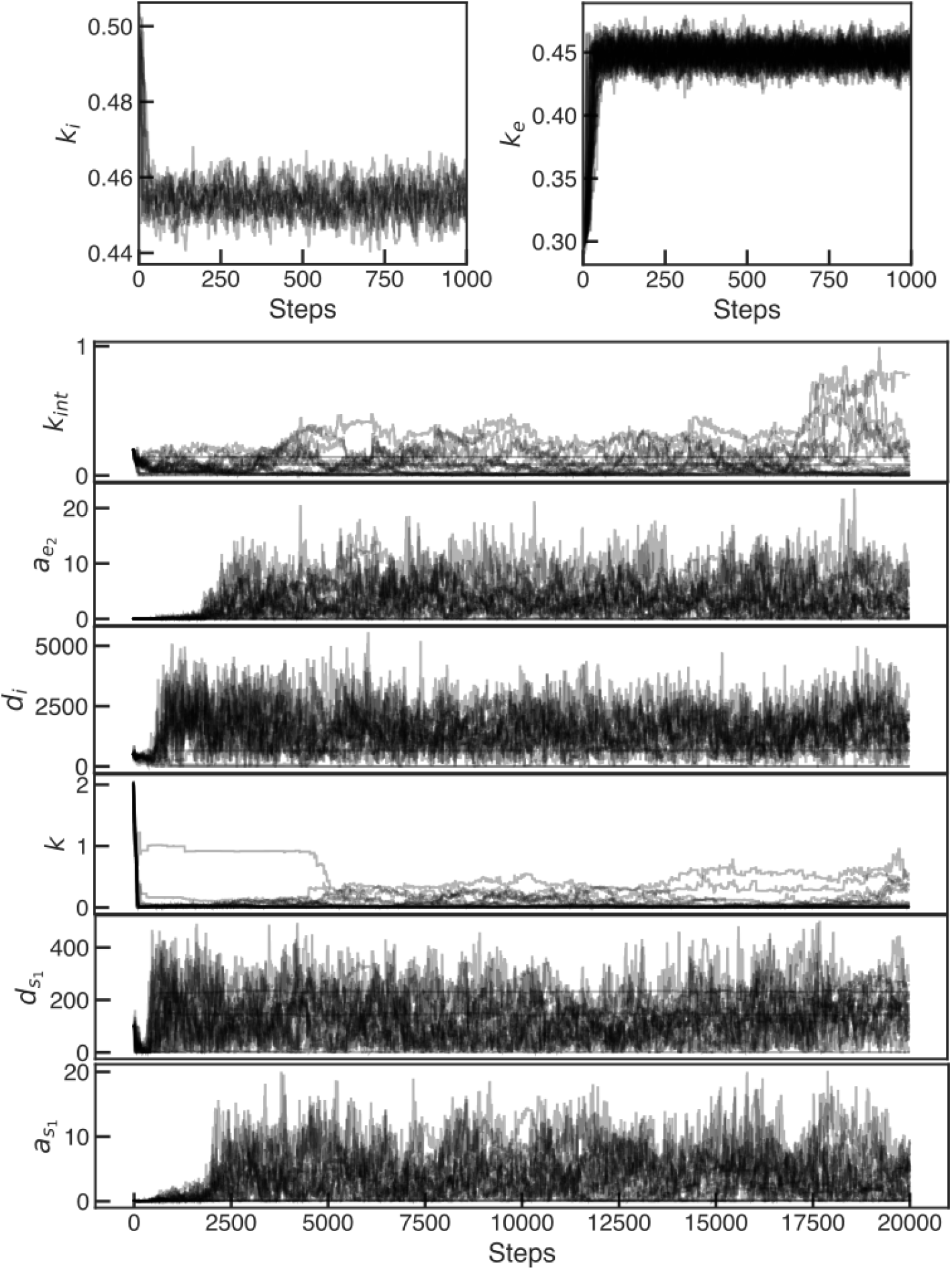
MCMC chains for identification of integrase-excisionase model parameters using the data from cell-free experiments.

#### Identified Excisionase Model and Data

We sample from posterior parameter distributions and run model simulations. We plot the simulations of the identified models together with the experimental data. These results are shown for all three measurements: CFP, mScarlet, and YFP in Figures 16, 17, and 18 respectively. As it is clear from the runtime, an increase in the dimension of the parameter inference problem from 3 to 6 led to a significant increase in runtime from around 1 hour to 24 hours. This is an expected curse of dimensionality, which leads to difficulty in inferring the parameters of detailed models. In the model-data fits shown above, the CFP and mScarlet measurements agree with the fitted model simulations, but, the YFP measurements do not perfectly fit the model predictions. The qualitative trends of the YFP expression are predicted correctly but since we are only identifying 6 parameters out of a total of 54 in the detailed model, these inaccuracies are expected.

**Figure 16:**
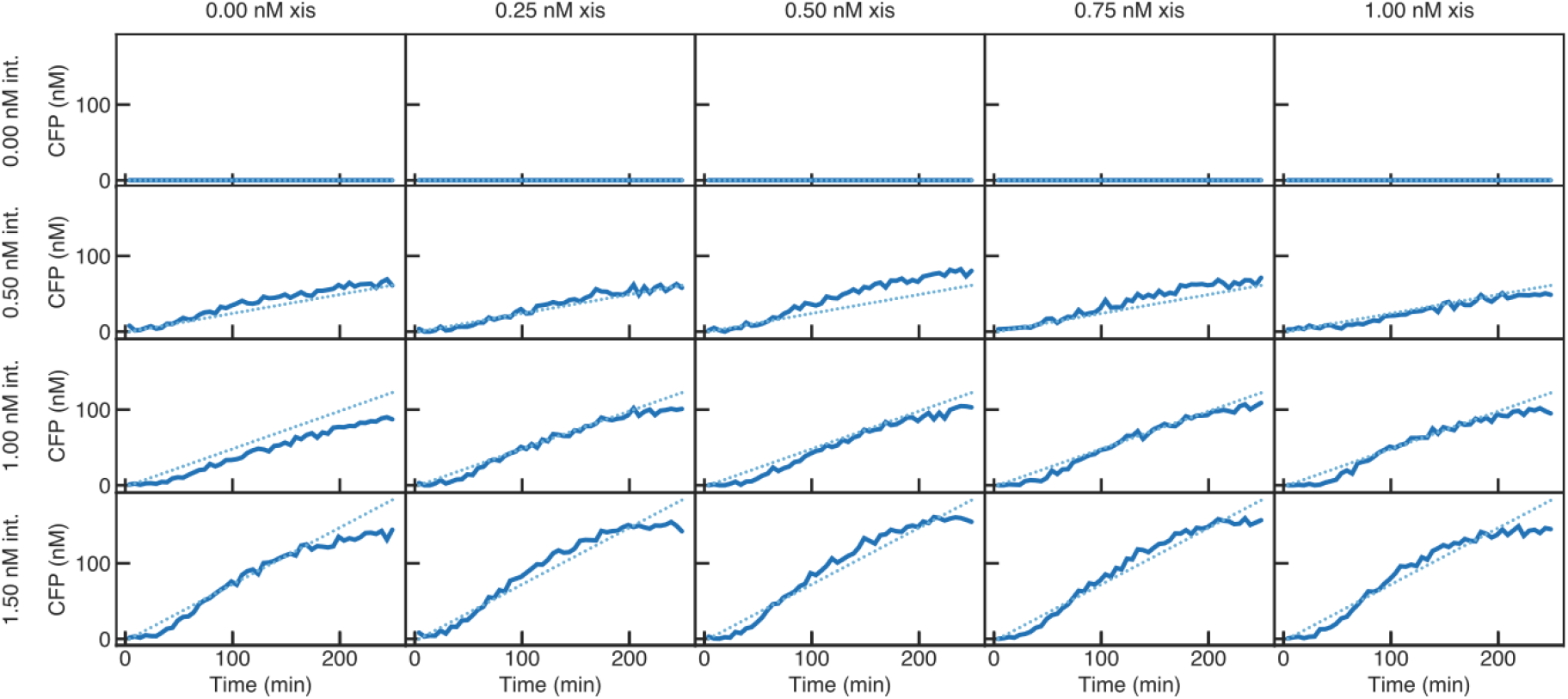
Model simulations with posterior parameter values plotted alongside CFP measurement.

**Figure 17:**
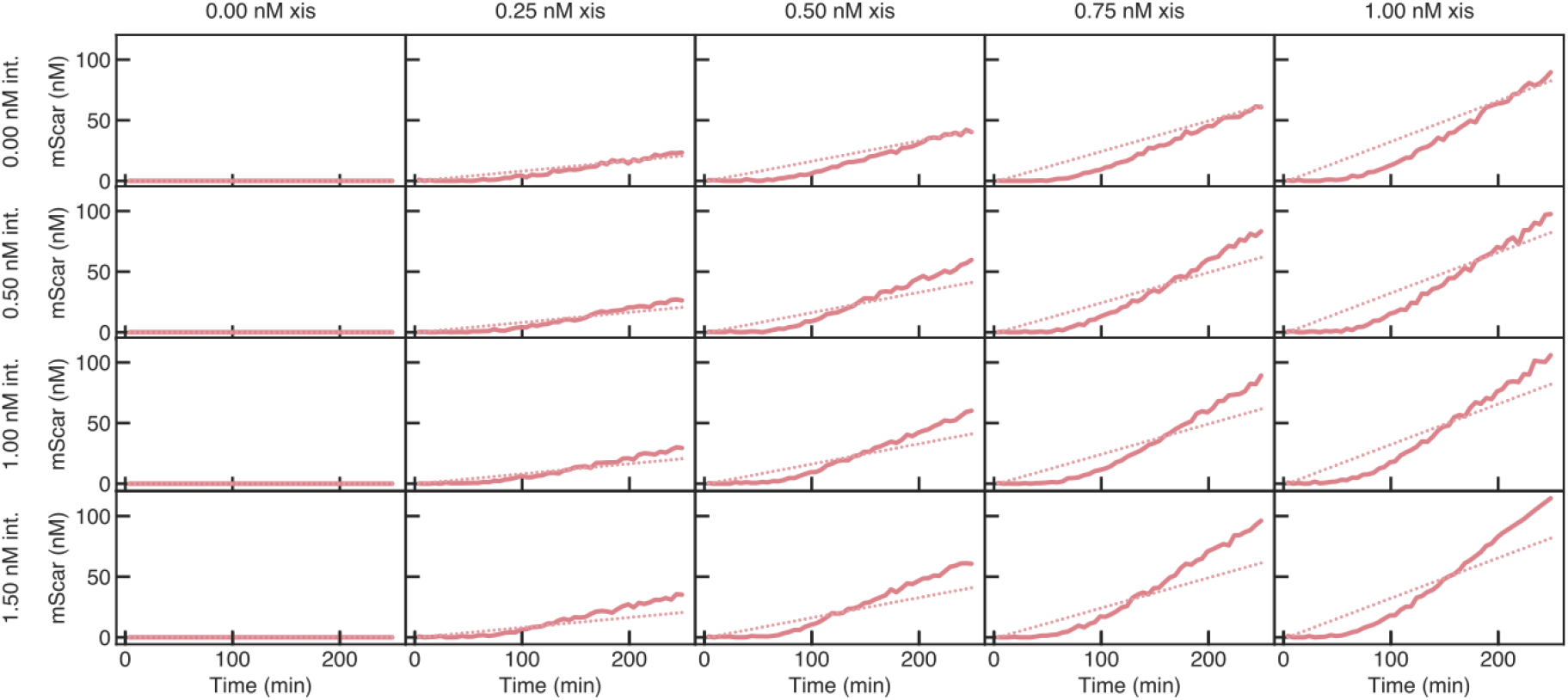
Model simulations with posterior parameter values plotted alongside mScarlet measurement.

**Figure 18:**
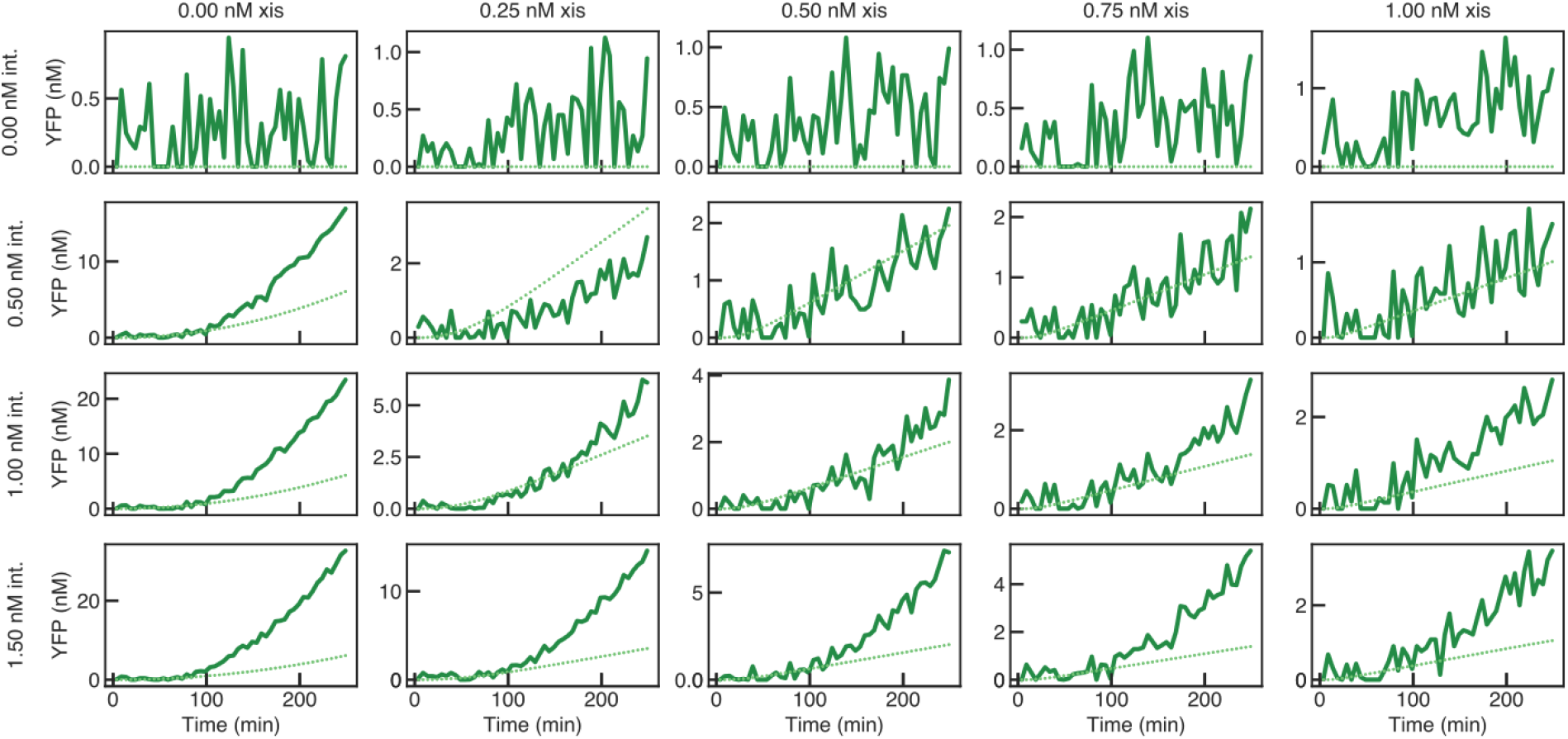
Model simulations with posterior parameter values plotted alongside YFP measurement. Note that although the model qualitatively captures the YFP expression but does not fit the data very well for some of the conditions. This is a result of a computational tradeoff in choosing a lower dimensional model for feasible parameter identification. The detailed model would capture the experimental behavior but it is infeasible to estimate 54 parameters with Bayesian inference.

#### Plasmid Designs

The integrase plasmid is composed of a pTet promoter^41^ followed by a strong BCD2 ribosome binding site.^42^ Downstream of the ribosome binding site is a fusion protein: Bxb1 integrase^43^ linked to a cyan fluorescent protein. Termination of transcription is achieved using a T500 terminator.^44^ The reporter plasmid contains a strong P7 promoter^42^ flanked by Bxb1 recombination sites attB and attP. Downstream of the attP recognition site is a RiboJ insulator^45^ and a BCD2 ribosome binding site^42^ driving the expression of a “Venus” yellow fluorescent protein.^46^ The integrase and reporter plasmids were taken from Swaminathan et al.^16^

The excisionase plasmid contains a pTet promoter^41^ directly followed by a strong BCD2 ribosome binding site.^42^ The promoter and ribosome binding site activate the transcription of a Bxb1 excisionase^43^ linked to a red fluorescent protein with a GS linker (a 12 amino acid sequence with 6 Glycine alternating with 6 Serine amino acids). Transcription is terminated using a T500 terminator.^44^

## Notes

https://github.com/BuildACell/BioCRNPyler

https://github.com/ayush9pandey/autoReduce

https://github.com/biocircuits/bioscrape/

https://github.com/ayush9pandey/integrase-excisionase-characterization

### Competing Interest Statement

The authors have declared no competing interest.

